# Functional genomics identifies therapeutic options, biomarkers, and resistance mechanisms for high-grade gliomas

**DOI:** 10.64898/2026.02.01.701806

**Authors:** Wan-Hsin Lin, Farhad Kosari, James B. Smadbeck, Michael T. Barrett, Ryan W. Feathers, Jamie Hall, Dorsay Sadeghian, Sotiris Sotiriou, Sarah H. Johnson, Faye R. Harris, Taylor Berry, Alexa F. McCune, Stephen J. Murphy, Lindsey Kinsella, Lauren E. Haydu, Diogo Moniz-Garcia, Shannon P. Fortin Ensign, Lin Yang, Angela R. Emanuel, Leila A. Jones, Janet L. Schaefer-Klein, Cristiane M. Ida, Marcela A. Salomao, Wendy J. Sherman, Alyx B. Porter, Steven S. Rosenfeld, Sani H. Kizilbash, Kurt A. Jaeckle, Maciej M. Mrugala, Aaron S. Mansfield, Mitesh J. Borad, Bernard R. Bendok, Terry C. Burns, Alfredo Quinones-Hinojosa, John C. Cheville, George Vasmatzis, Panos Z. Anastasiadis

## Abstract

High-grade gliomas (HGGs) are aggressive tumors with poor outcomes and limited treatment options. Here, we combined genomic and transcriptomic tumor profiling with drug testing in a patient-derived 3-dimensional culture model to identify individualized treatments and predictive biomarkers. Activity of single agents targeting frequently dysregulated glioma pathways was relatively poor *ex vivo* and generally reflected historical patient data. However, compounds targeting PI3K, epigenetic, and survival/senescence signaling were effective in some cases. Drug sensitivity correlated with transcriptional rather than genomic features and suggested heterogeneity as a resistance mechanism. Bromodomain and extraterminal domain inhibition was particularly effective in tumors enriched in the mesenchymal transcriptional subtype, promoted proneural transition, and was overcome by upregulated PI3K signaling. Notably, combinations were largely effective, with 6 strategies exhibiting stronger efficacy than corresponding single agents in most cases (58-77%). This study identifies HGG vulnerabilities and associated biomarkers, resistance mechanisms, and effective combination strategies that warrant further clinical validation.

**Graphical Abstract:** 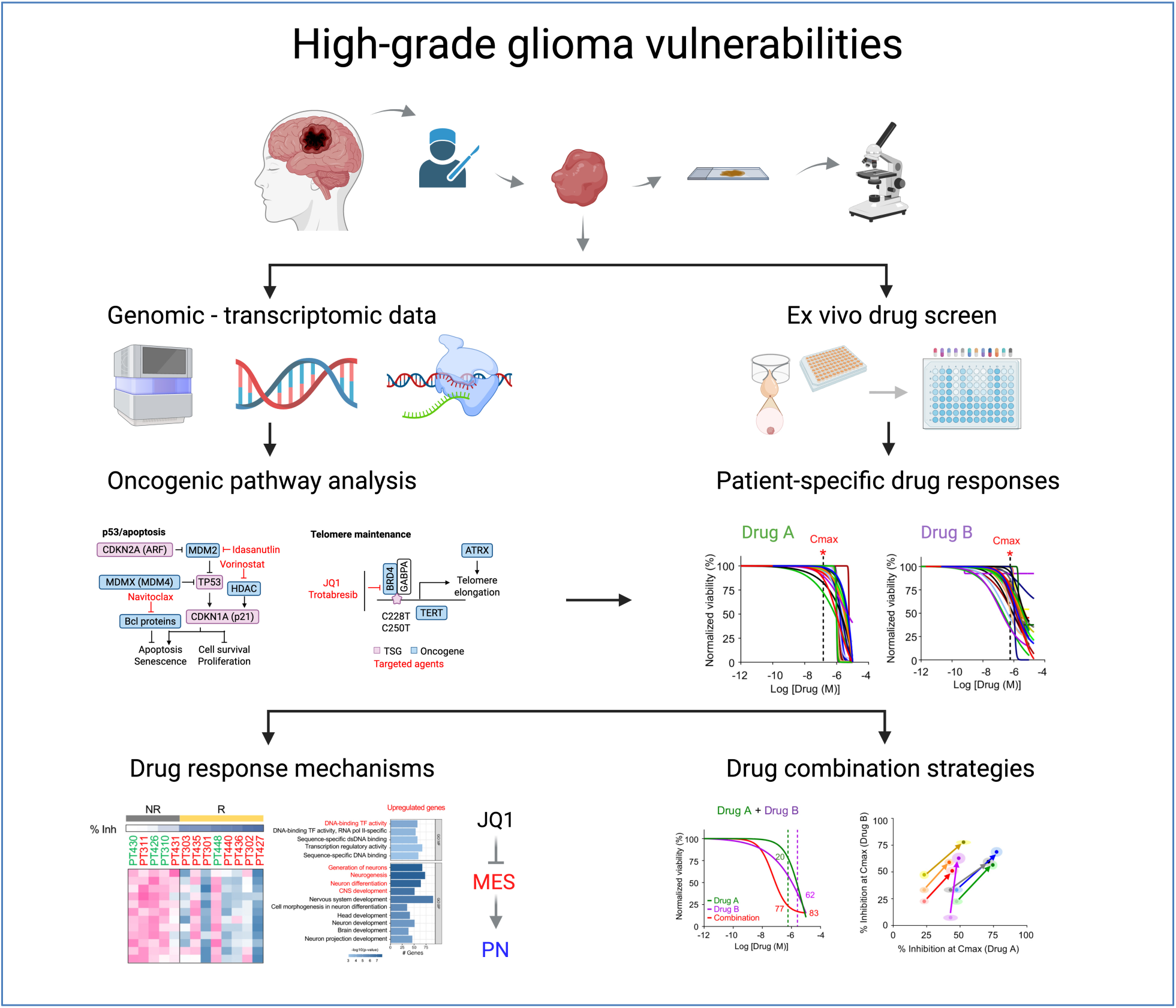

## Introduction

Diffuse gliomas are the most common malignant brain tumors in adults and include isocitrate dehydrogenase mutant (IDH-mut) astrocytomas, IDH-mut and chromosome 1p/19q-codeleted oligodendrogliomas, and IDH-wildtype glioblastomas (GBMs) (1). Tumor profiling has shown that high-grade gliomas (HGGs) are molecularly complex, with multiple cancer-driving events and oncogenic pathways concurrently deregulated (2). Possibly due to this complexity, clinical trials targeting specific alterations and/or designed based on drug efficacy in existing pre-clinical models have generally failed (3), underscoring the urgent need for novel approaches.

Conventional glioma cell lines (*in vitro* and *in vivo*) and orthotopic patient-derived xenografts (PDXs) have been extensively used in pre-clinical studies. More recently, stem cell-enriched neurospheroids and matrix-containing patient-derived organoids (PDOs) (4–7) have been used to interrogate therapeutic vulnerabilities. However, the time required for the generation of these models invariably introduces selective pressure due to culture conditions or murine-specific evolution (8), resulting in loss of cellular and molecular heterogeneity as well as key HGG features that are crucial for the translation of pre-clinical findings to patient treatment (9,10).

Here, we used a functional genomics approach that incorporates molecular profiling and a patient-derived 3-dimensional (3D) culture model we termed microcancer (μCancer) to enable rapid interrogation of HGG vulnerabilities to targeted treatments. This short-term culture model maintained heterogeneity and recapitulated the genomic and transcriptomic landscapes of parental patient tumors. Several treatment strategies, including compounds targeting the PI3K pathway, epigenetic signaling, and senescence were effective in select cases, with transcriptional features showing better predictive potential than genomic alterations. Bromodomain and extraterminal domain (BET) inhibition was particularly effective in tumors enriched in the mesenchymal transcriptional subtype, promoted proneural transition, and was overcome by upregulated PI3K signaling. Finally, six combination treatment strategies with stronger efficacy (58-77% of cases tested) than single agents and acting across glioma subtypes were also identified and warrant further clinical validation.

## Results

### ExVivo-Glioma functional genomic workflow

To identify potential therapeutic vulnerabilities and predictive biomarkers, we developed a functional genomics workflow called ‘*ExVivo-Glioma’* (Figure 1A). In this platform, resected tumor tissues were examined by a neuropathologist and divided into two, with one half immediately flash-frozen for subsequent genomic and transcriptomic profiling and the other half cryopreserved for μCancer generation and drug testing.

**Figure 1.**
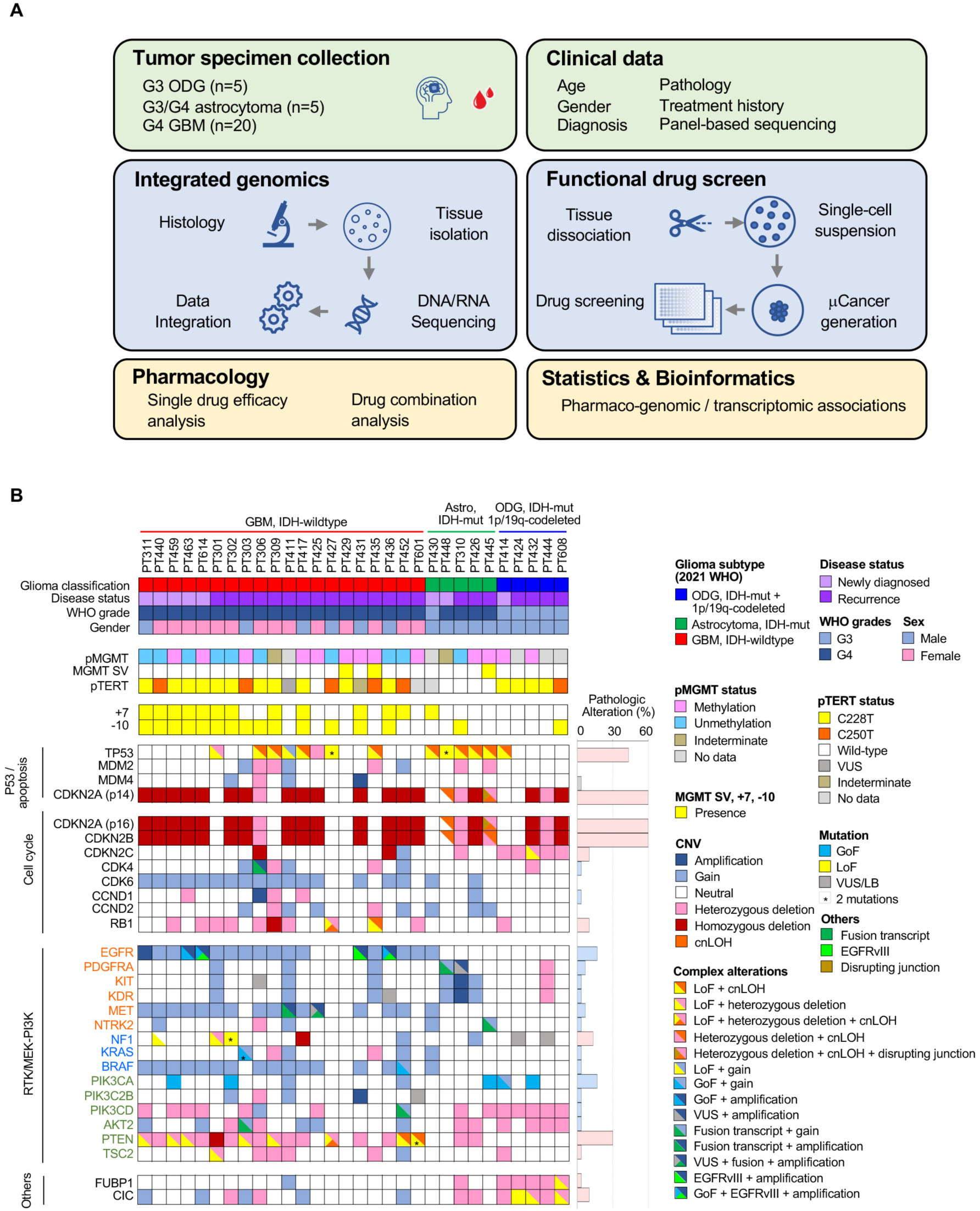
**Functional ExVivo-glioma platform to identify individualized targeted treatments for HGG.** (A) Overview of the ExVivo-glioma platform (created with BioRender.com). (B) Clinical parameters and glioma relevant molecular alterations from tumor tissue. RTK (orange), MEK (blue), and PI3K pathway (green) components indicated. Histogram (right) depicting the frequency of pathogenic alterations of listed genes (tumor suppressor genes (TSGs, pink) exhibiting or loss-of-function (LoF) mutation(s) or complete inactivation; oncogenes (blue) exhibiting amplification, fusion, or gain-of-function (GoF) mutation(s)). Notes: 2021 WHO classification used for disease diagnosis. PT601: hypermutated. LB, likely benign. See also Tables S2 and S3.

### Study population and clinical characteristics

A total of 47 glioma cases were initially enrolled with 9 being excluded due to patient withdrawal or lack of adequate tissue (Figure S1). Eight cases failed to establish μCancers or did not pass viability quality control assessments, for an overall μCancer success rate was 78.9%. The remaining 30 cases include twenty grade 4 GBMs, five grade 3 or grade 4 IDH-mut astrocytomas, and five grade 3 oligodendrogliomas (Figure 1A). One case (PT448) was complex with a low-grade IDH-mut component and a high-grade component that had lost the IDH-mut. Additionally, our study focused primarily on the recurrent setting (*n*=22) with inclusion of some newly diagnosed cases (*n*=8) (Table S1). Patients with recurrent disease received 1-5 prior treatments including chemoradiation, bevacizumab, and tumor-treating fields (Tables S1 and S2). Our cohort included more females (*n*=14) than males (*n*=6) in GBM and exclusively males in astrocytomas and oligodendrogliomas (Figure 1B; Table S1). Gliomas were located in the supratentorial region of the brain, including the frontal lobe (*n*=12), parietal lobe (*n*=3), temporal lobe (*n*=5), occipital lobe (*n*=1), or with involvement of two different lobes (*n*=9) (Table S2). *MGMT* promoter methylation in GBMs was present in 9 cases, absent in 9, and not tested or indeterminate in 2 cases (Figure 1B; Tables S1 and S2).

### Genomic and transcriptomic characterization

Pathology review of frozen tumor sections was first performed before DNA and RNA isolation. Mate-pair sequencing (MPseq) was used to produce whole-genome profiles to identify structural aberrations, including DNA rearrangements, copy number alterations (CNAs), and loss of heterozygosity (LOH). Somatic and germline whole exome sequencing (WES) were performed to identify mutations. Gene expression was investigated by RNAseq, while gene fusions were investigated by RNAseq and MPseq (11).

Most GBMs (55%) in our cohort exhibited the combined gain of the entire chromosome 7 (+7) and loss of the entire chromosome 10 (−10) (Figure 1B). Examining core signaling pathways that are frequently deregulated in glioma (2), we observed dysfunction of the p53/apoptosis pathway in 90% of cases (Figure 1B; Table S3), with *TP53* biallelic inactivation observed in 35% of GBMs and all IDH-mut astrocytomas. Further, our cohort contained 3 cases with low-level copy number gains of *MDM2*, and 1 containing co-amplification of *MDM4* and *EGFR*.

Biallelic inactivation of *CDKN2A* and *CDKN2B* were the most frequent alterations related to cell cycle signaling, present in 70% of GBMs and 30% of IDH-mut gliomas (Figure 1B; Table S3). Amplification of *CCND1* was observed in one case, while complete *RB1* inactivation through various alterations was found in three cases. Combined, deregulation of cell cycle signaling was observed in 73% of cases.

*EGFR* amplification was found in 5 cases with one expressing wild-type EGFR, one expressing an ectodomain p.A289D mutation, and the remaining expressing *EGFRvIII* variant (Figure 1B; Table S3). Two *EGFRvIII*+ cases concurrently exhibited ectodomain mutations. Further, our cohort included 2 amplified *PTPRZ1::MET* fusion GBMs and 2 *PDGFRA*-positive astrocytomas, one amplified and one expressing a *LNX1::PDGFRA* fusion transcript. Downstream of receptor tyrosine kinases (RTKs), 3 cases exhibited complete *NF1* inactivation, and 1 expressed an oncogenic *KRAS* mutation. In general, genomic alterations related to PI3K signaling occurred more frequently (47%) than MEK pathway alterations (20%), with complete inactivation of *PTEN* being the most common alteration. Overall, genomic alterations in RTK/MEK/PI3K signaling were observed in 73% of cases.

### Patient-derived μCancers retain the molecular landscapes and salient features of parental tumors

To minimize tumor evolution, subclonal selection, and loss of genomic and cellular heterogeneity, we developed a short-term 3D culture model that can be rapidly established from fresh or cryopreserved tumor tissues (Figure 2A) (11). Briefly, bulk tumor was enzymatically and mechanically dissociated into single cells using a gentleMACS^TM^ Dissociator. Dissociated cells were then plated onto 96-well Akura^TM^ PLUS hanging-drop plates in optimized culture media and allowed to re-aggregate for 6 days to form 100-300 μm-sized 3D μCancers, as before (11). Patient-derived μCancers retained a similar Ki-67 proliferation index as their parental tumors and maintained proliferative capacity, exhibiting a cell doubling time of ∼7 days (Figure S2A-C). This rate of growth suggests that both cytotoxic and cytostatic effects can be interrogated in μCancers over a 6-day drug treatment, and that short-term culture would minimize selective pressure, thus maintaining the genomic profile of the original tumor.

**Figure 2.**
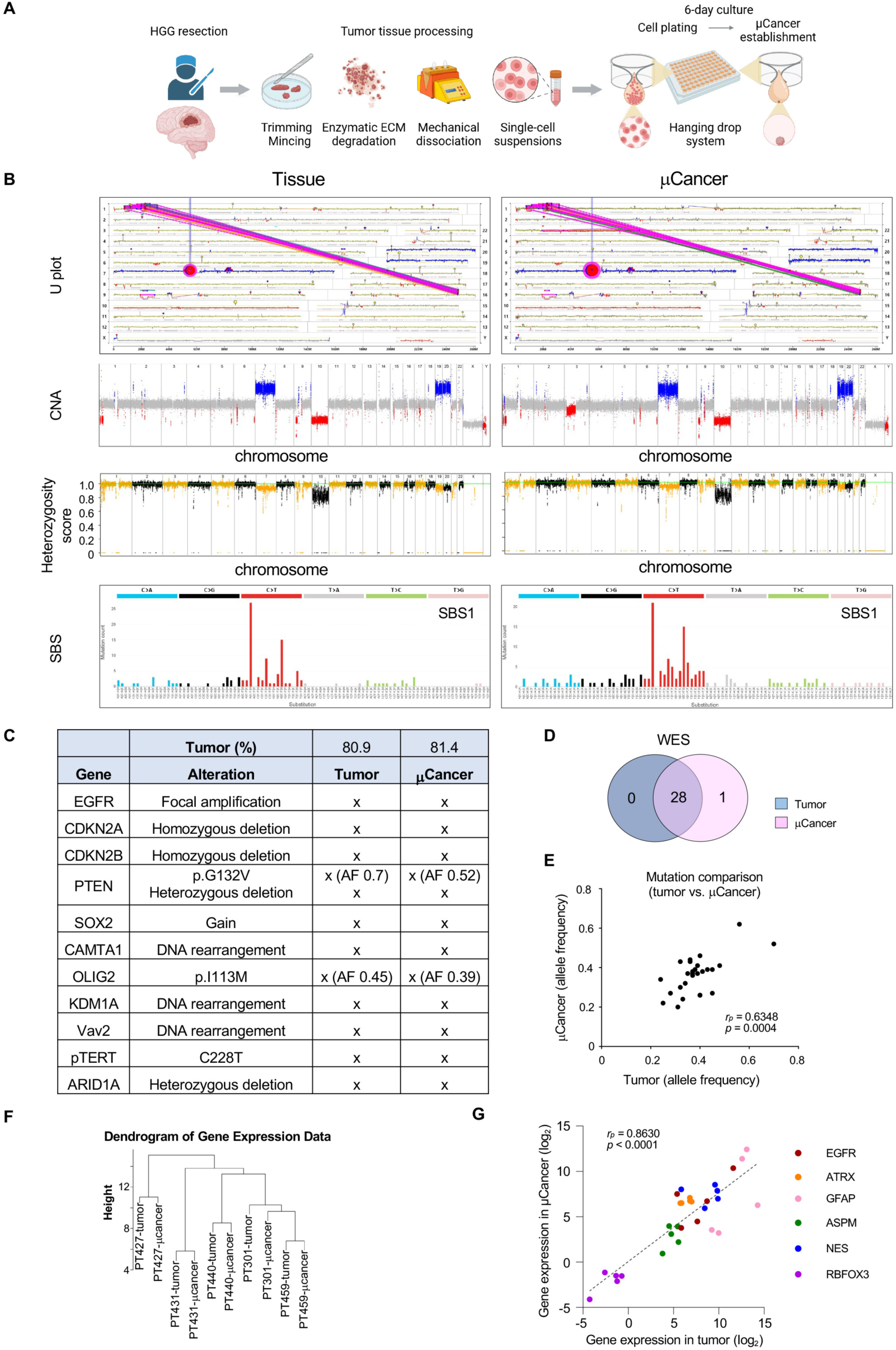
**Comparison of patient-derived μCancers and parental GBM.** (A) Workflow of μCancer establishment from resected HGG tissue (created with BioRender.com). (B) Genomic landscape showing CNAs and mutations (lollipops) in U plot (top), horizontal CNAs (middle top), heterozygosity score (middle bottom), and SBS signature (bottom). Gains (blue), losses (red), diploid (grey), and DNA junctions (magenta) indicated. (C) Table showing glioma-relevant mutations identified in parental tumor and retained in μCancers. Allele frequency (AF) indicated. (D) Venn diagram illustrating mutations from WES overlapped between μCancer (pink) and parental tumor (blue). (E) Correlation plot showing comparable allele frequency of mutations from WES in μCancer and parental tumor. (F) Dendrogram depicting clustering patterns of 5 μCancer – control tumor pairs. (G) Correlation plot showing expression of indicated marker genes between tumor and μCancer samples from the five matched pairs in E. Pearson’s correlation (*r_p_*) and *p*-value indicated. See also Table S4.

To test whether μCancers maintain the genomic profile of parental tumors, MPseq and WES were performed on PT311-derived μCancers cultured for 6 days (Figures 2B-2E). μCancers faithfully retained the +7/-10 signature as well as the focal amplification of *EGFR* on extrachromosomal DNA (Figure 2B, top two panels), an event frequently lost during cell culture (12,13). Further, parental tumor and resulting μCancers exhibited similar heterozygosity and the same single base substitution signature SBS1 (Figure 2B, bottom two panels), as well as key oncogenic events and tumor percentage (Figure 2C). μCancers also preserved all 28 mutations identified in the original tumor with similar allele frequencies, indicating they maintain the genomic diversity of parental tumors (Figures 2D and 2E; Table S4). Whole-genome CNAs and DNA rearrangements (95 of 98 events; 96.9%) were also preserved (Figure S2D). Minor subclonal molecular alterations are likely due to the inherent spatial heterogeneity of these tumors, as sequencing for the parental tissue and corresponding μCancers was performed in adjacent but distinct segments of the same tumor (one flash-frozen and the other cryopreserved). Finally, the μCancer *TERT* promoter mutation status was identical to that of the parental tumor (Figure S2E).

Transcriptomic fidelity was assessed in five GBM cases, 2 primary (PT440, PT459) and 3 recurrent (PT301, PT427, PT431), and each tumor sample clustered or grouped in close proximity with its corresponding μCancer (Figure 2F). Using single-cell RNAseq, we previously reported that μCancers derived from GBM8 PDX tumors maintain the cellular diversity of the original tumor including the presence of glioma stem cells, differentiated glioma cells, and murine stroma (14). In agreement, the expression of glioma tumor markers (*EGFR*, *ATRX*), astrocytic lineage marker (*GFAP*), neurogenesis-related glioma stem cell regulators (*ASPM*, *NES*), and neuronal marker (*RBFOX3*), were highly correlated between original patient tumors and their resulting µCancers (Figure 2G). Overall, several lines of evidence indicate that μCancers recapitulate the genomic and transcriptomic landscapes and heterogeneity of parental tumors with high fidelity, providing an optimal testbed for the investigation of potential HGG therapies.

### Functional drug screen

Established μCancers, derived from dissociated tumor cells cultured for 6 days, were transferred to ultra-low attachment (ULA) plates to enable drug testing (Figure 3A). Initially, we compared the effect of different media formulations on cell viability and drug responses (Figure S3A). Overall metabolic activity, reflecting cell viability, was higher in the μCancer media, although responses to select targeted agents, including pimasertib, capivasertib, and navitoclax, were similar regardless of media formulations. A good correlation was also observed in drug responses between repeats of the same sample performed at different times (Figure S3B).

**Figure 3.**
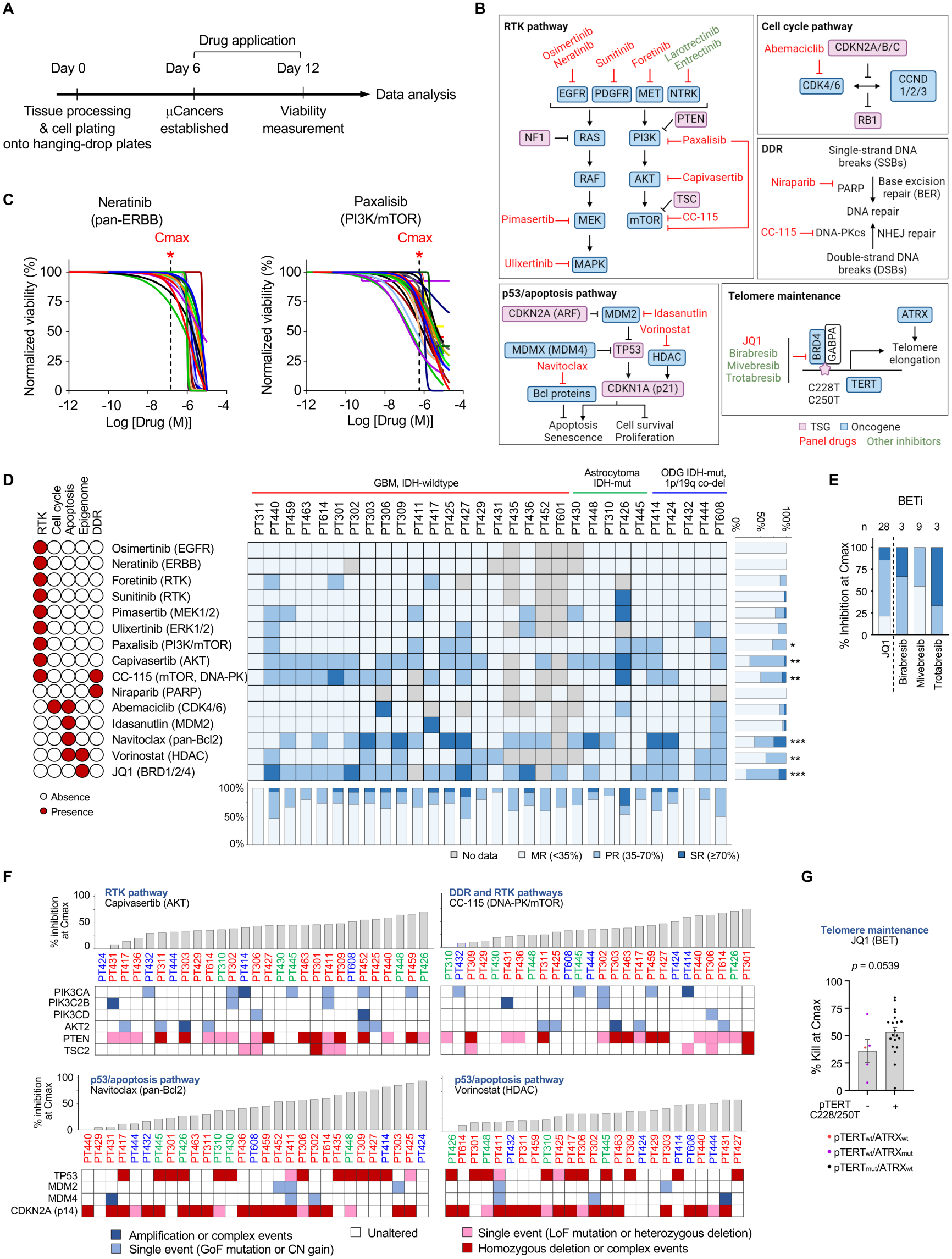
***Ex vivo* single-agent efficacy profiling and correlation with genetic alterations.** (A) Workflow for μCancer generation and drug testing followed by biomarker identification. (B) Schematic illustrating agents targeting key glioma pathways (created with BioRender.com). (C) Normalized μCancer dose-response curves for indicated agents with Cmax indicated (vertical dashed lines with asterisks). Each trace represents one case. (D) Three-tier drug response heatmap (middle) showing <35% (minimal response, MR; light blue), 35-70% (partial response, PR; blue), and ≥70% (strong response, SR; dark blue) of growth inhibition at Cmax for indicated inhibitors (left). Stacked bar graphs depicting percentage of differential responses for each drug (right) and case (bottom). \**p*<0.05, \*\**p*<0.01, \*\*\**p*<0.0005 vs. osimertinib (least responsive drug) by multivariable GLMM. (E) Stacked bar graph showing percentage of differential responses for BET inhibitors. (JQ1 from D plotted for comparison). (F) Waterfall plots of % inhibition at Cmax for μCancers treated with indicated agents (top). Drug target-related genetic features summarized (bottom; oncogenes and TSGs colored in blue and red shades, respectively). Case IDs colored by histotypes, as in panel D. (G) Scatter plot with bars (mean±SEM) showing JQ1 efficacy in μCancers from tumors with wildtype or mutated *pTERT*, also indicating cases with *ATRX* alterations. *P*-value determined by unpaired t-test (n=25). See also Tables S6 and S8.

A total of 116 single agents and 183 drug combinations were tested based on the presence of actionable alterations and included a pre-selected drug panel that targets frequently deregulated HGG pathways (2). This panel, which is the focus of this work, included 16 agents targeting deregulated RTK signaling, p53/apoptosis, cell cycle, DNA damage and repair (DDR), telomere maintenance mechanisms, and IDH signaling (Figures 3B and S4A), along with 7 drug combination strategies that will be discussed later. When possible, drugs in each class were selected based on their ability to penetrate the central nervous system (CNS) (Table S5). Alternatively, compounds targeting key pathways and/or approved for use in other clinical settings were chosen.

Following μCancer generation and drug testing, drug efficacy was represented by the percentage of reduced viability (% inhibition) at the maximum serum concentration of the drug achieved after dosing (Cmax) (see Figure 3C, Table S5). To facilitate result interpretation, we segregated response profiles into three categories: <35%, 35-70%, and ≥ 70% inhibition at Cmax that were defined as minimal response, partial response, and strong response, respectively (Figure 3D). Samples with sufficient tissue were subjected to testing with all the drugs in the panel (Table S6).

### Limited vulnerability of HGGs to single agent treatments *ex vivo*

Glioma μCancers showed in general minimal response to RTK-targeted agents, including the EGFR inhibitor osimertinib, pan-ERBB inhibitor neratinib, and multi-RTK inhibitors foretinib and sunitinib (Figures 3B-3D and S3C; Table S6). Similar results were also obtained by testing select cases with additional EGFR-related inhibitors, alone or in combination (Tables S6 and S7). Additionally, the efficacy of foretinib or sunitinib treatment was unrelated to molecular alterations of their targets MET and KDR, or KDR, PDGFRA, and KIT, respectively (Figures 1B and 3D).

To inhibit RTK signaling by targeting downstream effectors, we utilized the MEK inhibitor pimasertib, MAPK/ERK inhibitor ulixertinib, PI3K/mTOR dual inhibitor paxalisib, pan-AKT inhibitor capivasertib, and mTOR/DNA-PK dual inhibitor CC-115. In general, this strategy was more effective than targeting upstream RTKs (Figure 3D). Inhibitors targeting the PI3K pathway were significantly more effective than osimertinib, the least effective agent tested (Figure 3D and Table S8). Paxalisib achieved 8 partial responses; capivasertib, 1 strong and 19 partial responses; and CC-115, 2 strong and 11 partial responses. The extent to which the DNA-PK (DDR kinase) inhibitor activity of CC-115 contributes significantly to these responses is currently unclear, although another DDR-targeting agent, the PARP inhibitor niraparib, exhibited no effect in our μCancer cohort.

Despite common activation of the cell cycle pathway in gliomas via *CDKN2A/B* homozygous deletion (Figure 3B), μCancer treatment with the CDK4/6 inhibitor abemaciclib was largely ineffective (Figure 3D). Notably PT306, which responded strongly to abemaciclib, exhibited focal amplifications of *CCND1* and *CDK4::MYEOV* fusion, a potential oncogenic driver (Figure 1B; Table S3). In agreement with its low efficacy in μCancers, abemaciclib was largely ineffective in patients with recurrent GBM (Table S5).

Idasanutlin, an MDM2 inhibitor used to restore p53 pathway activation (Figure 3B), had limited effect in μCancers (Figure 3D), consistent with the lack of *MDM2* amplification in our cohort, which was associated with response to MDM2 inhibitors in PDX models (15). Moreover, amplification of *MDM4* in PT431 did not confer response to idasanutlin. In contrast, the pan-Bcl-2 inhibitor navitoclax was effective, exhibiting 7 strong and 11 partial responses *ex vivo* (Figure 3D). Interestingly, navitoclax can eliminate senescent GBM cells that are normally resistant to apoptosis (16). The data argue that agents that restore apoptosis and also act as senolytics may have strong activity in glioma, a hypothesis that warrants further investigation.

Multiple studies argue that the epigenome is largely altered during gliomagenesis and disease progression (17,18). Here, we targeted the epigenome by testing the activity of histone deacetylase (HDAC) and BET inhibitors, both shown to modulate chromatin structure organization to globally affect gene expression (19). HDAC inhibition by vorinostat resulted in moderate but statistically significant activity, while JQ1, a preclinical BET inhibitor, was significantly more effective (Figure 3D and Tables S6 and S8). Similar responses were observed when 3 clinically relevant BET inhibitors, birabresib, mivebresib, and trotabresib (the latter been actively explored in glioma clinical trials) (20) were tested in select cases (Figure 3E; Table S6). These inhibitors are particularly active against the BET protein BRD4 and can displace it from chromatin leading to anti-tumor activity via gene expression modulation (19).

The IDH-mut inhibitor vorasidenib improved progression-free survival (PFS) and was approved for low-grade glioma patients with no prior treatments except surgery (21) (Table S5). However, no evidence to date supports its efficacy in the HGG setting (22), where we observed no effect on the viability of IDH-mut-derived μCancers (Figure S4B). All IDH-mut tumors in our cohort were high-grade and most of them (7 of 10) were pretreated recurrent cases (Tables S1 and S2).

Overall, although most of the panel drugs had limited single-agent activity, we observed at least a partial response in over 40% of cases for five agents, namely navitoclax, JQ1, capivasertib, CC-115, and vorinostat. Moreover, 46.7% of cases responded strongly (i.e., ≥70% inhibition at Cmax) to at least one of the 16 agents in the panel (Figure 3D; Table S6).

### Drug efficacy correlates better with transcriptomic rather than genomic HGG alterations

Next, we explored whether the effects of these five most effective single agents were associated with the presence of specific genomic alterations. Drug response to capivasertib and CC-115 showed no correlation with genomic alterations in *PIK3Cs*, *AKT2*, *PTEN*, or *TSC2* (Figure 3F). For navitoclax, we compared treatment response with oncogenic alterations in the p53/apoptosis pathway and found no direct correlation. No correlation was also observed between genomic alterations in the p53/apoptosis pathway and μCancer responses to vorinostat, tested here because HDAC inhibition can increase p53 stability and activate p21-mediated apoptosis in a p53-independent manner (23).

Effects of epigenetic drugs (epi-drugs) are likely attributed to overall changes in gene expression, through their action on chromatin and super-enhancer occupancy (19). However, BRD4 inhibition by JQ1 also regulates telomere maintenance (Figure 3B) by decreasing TERT expression in cells with *pTERT* mutations, including GBM stem cells (24), and by suppressing telomere elongation (25). In agreement, cases with *pTERT* mutation tended to respond better to JQ1, without reaching statistical significance in our cohort (Figure 3G).

We next investigated whether the overall burden of genomic alterations within drug-targeted pathways correlates with treatment response. A bioinformatics metric termed ‘differential ploidy’ was used to evaluate CNAs across all genes in the pathway that deviated from tumor ploidy. This method accounts for oncogenes and tumor suppressor genes (TSGs) and provides an additional measure of pathway activation. Focusing on well-established KEGG genes within each targeted pathway (Table S9), we found no correlation between pathway-specific differential ploidies and µCancer response to capivasertib, navitoclax, and CC-115 (Figure 4A).

**Figure 4.**
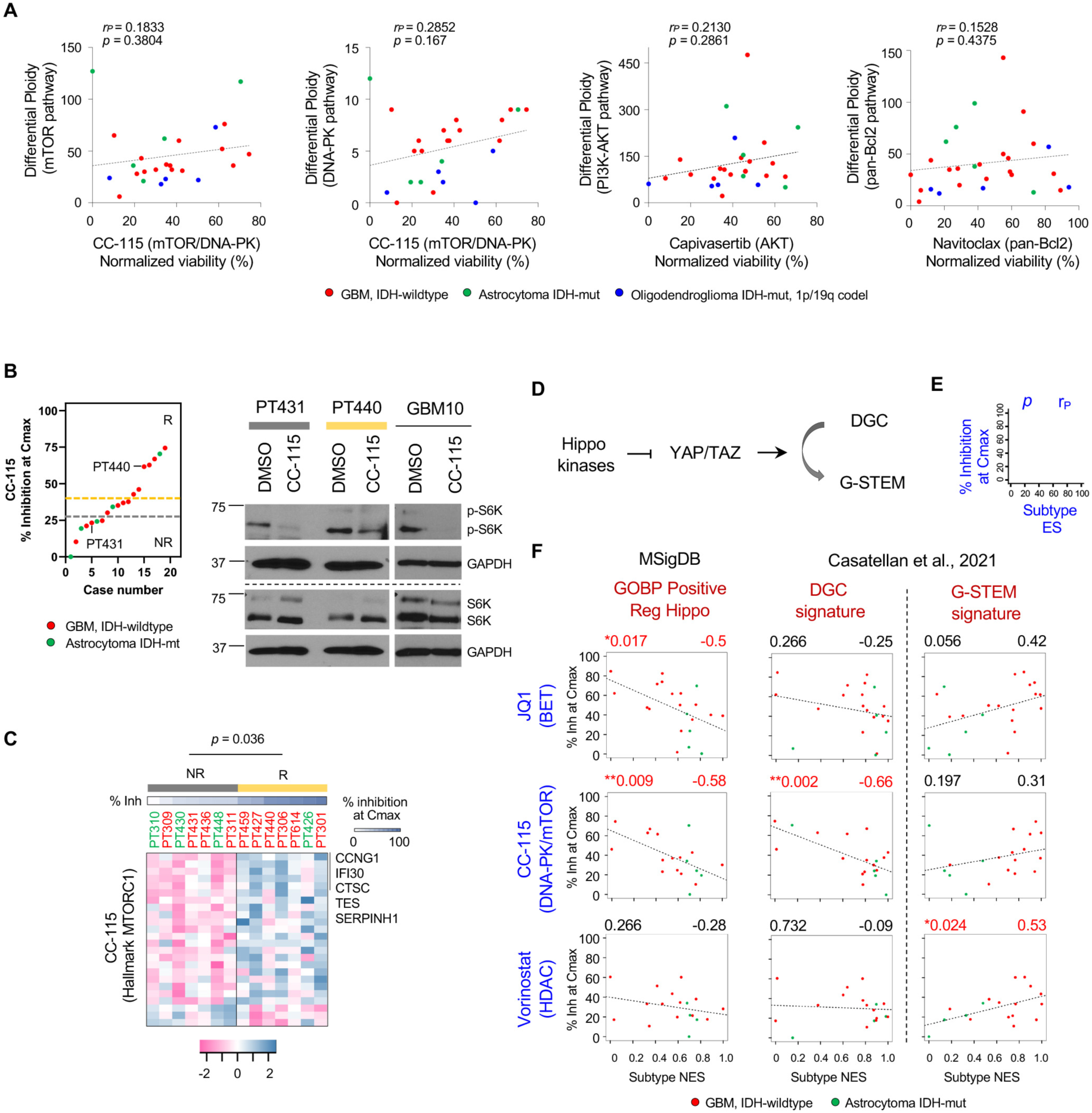
**Transcriptional signatures of Hippo pathway inactivation, YAP/TAZ activation, and glioma stemness linked to μCancer drug response.** (A) Correlation plots showing the relationship between µCancer response and differential ploidy of drug-targeted pathways. Pearson’s correlation coefficient (r_p_) with *p*-values and linear regression (dashed lines) indicated. (B) Distribution plot showing % inhibition at Cmax for cases ordered by increasing response. Responders (R) and non-responders (NR), above the yellow and below the grey lines, respectively with non-responding PT431 and responding PT440 indicated (left). Western blots showing phosphorylated and total S6K and GAPDH control in µCancers treated with CC-115 or DMSO (GBM10, 6-day treatment; PT431 and PT440, 2-day treatment). (C) Heatmap depicting MTORC1 hallmark genes differentially expressed between CC-115 responder (R; yellow) and non-responder (NR; gray). Z-scores (dark magenta to dark cerulean), % inhibition at Cmax (% Inh), and Pearson’s *p*-value indicated. (D) Schematic depicting Hippo kinases inactivation of YAP/TAZ signaling, preventing DGC-to-G-STEM conversion. (E) Graph key showing *p*-value (upper left) and Pearson’s correlation coefficient (r_p_, upper right) for panel F. (F) Correlation plots showing relationships between enrichment of indicated signatures (x-axis) and μCancer response to indicated agents (y-axis) with linear fits indicated. Hippo inactivation (GOBP Positive Reg Hippo, DGC) and YAP/TAZ activation (G-STEM) signatures separated by vertical line. Red denoted significant *p*-values and corresponding r_p_. See also Tables S9 and S10.

To assess whether the lack of correlation between drug response and genomic alterations is due to insufficient engagement of a drug’s target, we examined the effects of the dual mTOR/DNA-PK inhibitor CC-115 on the phosphorylation status of S6K, a key downstream effector of the mTORC1 complex (26). CC-115 treatment resulted in reduced phosphorylation of S6K in both non-responder PT431 and responder PT440, as well as GBM10 μCancers (Figures 4B and S5A-S5B). Consistent with its ability to target the mTORC1 complex, CC-115 response showed a significant positive correlation with a transcriptional signature for “genes up-regulated through activation of mTORC1 complex” (Hallmark mTORC1, Figure 4C; Table S10). Overall, the data suggest that although DNA alterations are key drivers of gliomagenesis, their combined effects on the transcriptome may better predict drug responses in µCancer models *ex vivo*.

### YAP/TAZ activation correlates with µCancer response to select single agents

Next, we postulated that the efficacy of JQ1, vorinostat, and CC-115 may be linked to their ability to modulate YAP/TAZ signaling, a key oncogenic pathway and therapeutic target in HGGs (27–30). JQ1 targets BRD4, a required cofactor that drives YAP/TAZ-mediated transcriptional addiction and tumorigenesis (31), whereas vorinostat targets HDACs, which facilitate transcriptional repression of TSGs by YAP/TAZ (32). CC-115 inhibits DNA-PK, which reportedly forms a functional complex with YAP/TAZ (33), and also suppresses mTOR activity, which is upregulated by YAP signaling (34). Inhibition of Hippo signaling promotes the transcriptional coactivator function of YAP/TAZ and drives tumorigenesis and progression via regulation of gene expression (Figure 4D) (35). Utilizing the “positive regulation of Hippo signaling” gene set from the Gene Ontology Biological Process (GOBP) class in the Molecular Signature Database (MSigDB), we observed that gliomas enriched for this Hippo activator gene set (corresponding to downregulation of YAP/TAZ signaling), were negatively correlated with μCancer response to JQ1 and CC-115 (Figures 4E and 4F). A negative trend between this gene set and response to vorinostat was also observed. The data suggest that YAP/TAZ activation confers μCancer sensitivity to JQ1 and CC-115, and possibly to vorinostat.

It is well-established that YAP/TAZ promote glioma stem cell (GSC) cellular states, hinder GSC transition to differentiation states, and promote resistance to treatment (27). To assess the role of GSC and differentiation states in μCancer responses to these agents, we compared each treatment response to the differentiated GBM cell (DGC) and G-STEM signatures (Figures 4D-4F) (27). We observed a negative correlation between enrichment for the DGC signature and μCancer response to JQ1 and CC-115, and a positive correlation between enrichment for the G-STEM signature and μCancer response to all three agents. These data are consistent with the hypothesis that YAP/TAZ signaling is an important target of these agents, regulating both glioma differentiated and stemness cellular states. Overall, these findings further argue that in molecularly complex tumors like HGGs, transcriptomic signatures correlate better with response to treatment than genomic alterations.

### HGG transcriptional subtypes predict µCancer drug responses

HGGs are characterized by high molecular complexity and cellular plasticity (36,37), which is reflected in the identification of several HGG transcriptomic cell states. The widely adopted The Cancer Genome Atlas (TCGA) classifier segregated gliomas into proneural (PN), classical (CL) and mesenchymal (MES) transcriptional subtypes (38). As YAP/TAZ is a master regulator driving MES characteristics in glioma (27), we next determined whether specific transcriptomic cell states correlate with μCancer drug responses. Our cohort included 15 PN, 11 CL, and 4 MES cases (Figure 5A). Further, to quantify transcriptional heterogeneity we applied a well-established ‘simplicity scoring’ (38) onto the classification. This scoring system ranges from 0 to 1, with 1 showing the simplest tumor that contains solely one subtype and 0 reflecting the coexistence of multiple subtypes within a single tumor. The average simplicity score (SPL) of our cohort was 0.50 ± 0.18 (mean±SD) with SPL of less than 0.5 observed in 14 out of 30 cases, indicating that multiple subtypes are present in many cases in our cohort.

**Figure 5.**
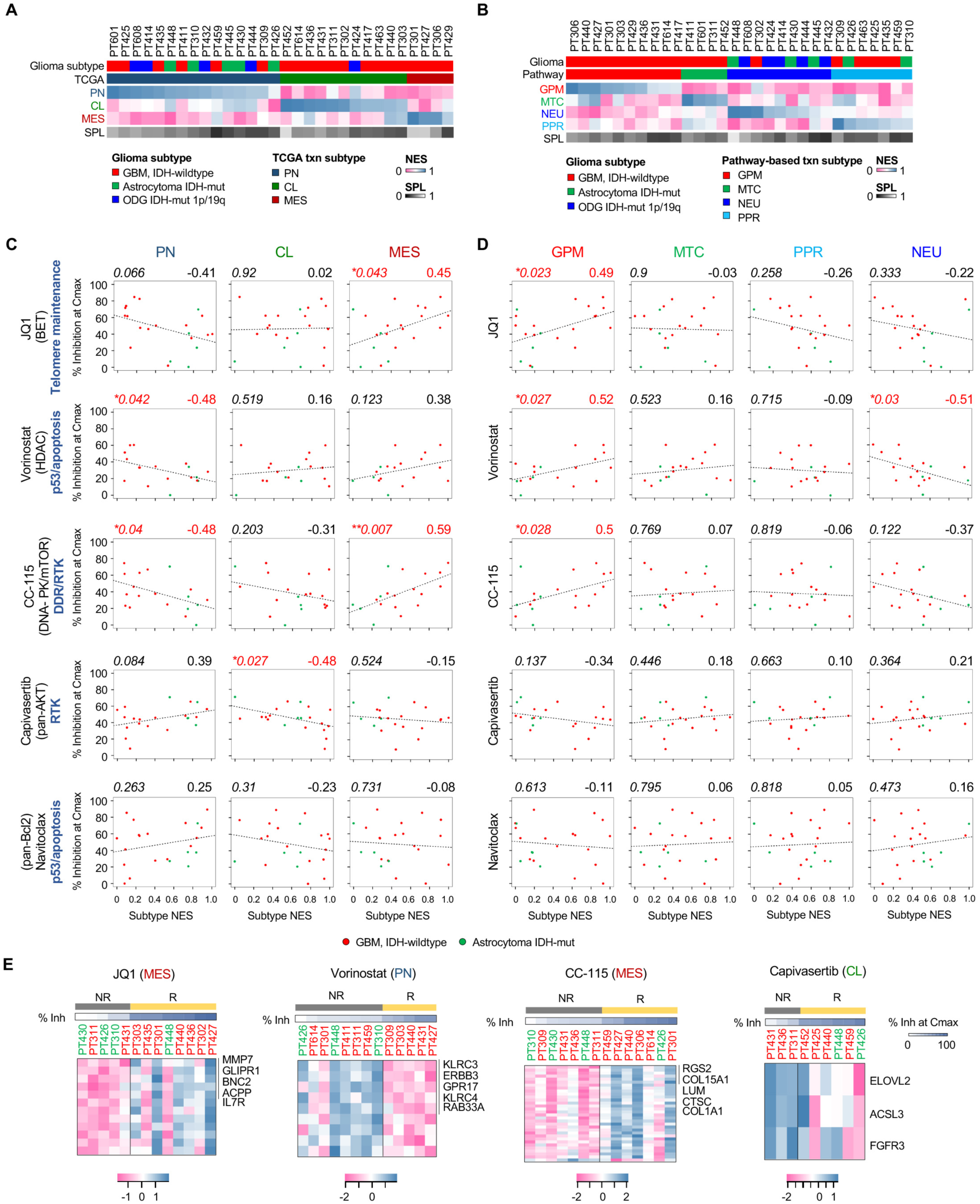
**Relationship between transcriptional subtypes and μCancer drug response.** (A-B) TCGA (A) and pathway-based (B) transcriptional subtypes (second row) with glioma subtype (first row) and normalized enrichment scores (NES) of each signature shown. NES indicating signature activation from high (1, dark cerulean) to none (0, dark magenta). SPL showing signature complexity from high (0, black) to low (1, white). (C-D) Correlation plots showing the relationship between normalized enrichment score (NES; scale: 0-1, x-axis) of indicated transcriptional subtype and μCancer response (% Inhibition at Cmax (% Inh); y-axis) to indicated agents. Drug targets (in parentheses) and associated pathways from Figure 3B (bold) indicated. *p*-value (upper left) and r_p_ (upper right) shown with red values denoting significance (\**p*<0.05, \*\**p*<0.01) with corresponding r_p_. GBM (red), astrocytoma (green), and linear regression lines indicated. (E) Heatmaps showing significant differentially expressed genes for indicated subtypes between responders (R, yellow) and non-responders (NR, gray) to indicated inhibitors. Z-scores from low (dark magenta) to high (dark cerulean) and % inhibition at Cmax (% Inh) indicated. Top 5 genes listed. See also Table S10.

A pathway-based transcriptomic classifier was recently developed for GBM subgrouping to predict biological activities underlying drug responses (39). According to this classification, our cohort consisted of 10 glycolytic/plurimetabolic (GPM), 4 mitochondrial (MTC), 9 neuronal (NEU), and 7 proliferative/progenitor (PPR) cases (Figure 5B), with MTC and GPM representing the best and worst prognostic subtypes, respectively (39). Further, SPL applied to this classification indicated that 19 of 30 cases exhibited SPL of less than 0.5. Comparing TCGA and pathway-based subtypes, all MES and half of CL gliomas in our cohort aligned with the GPM subtype, while PN tumors included MTC, NEU, and PPR subtypes, as previously reported (39) (Figures S5C and S5D).

To assess whether transcriptional subtypes provide predictive insight into response to μCancer treatment, we limited our analyses to GBM and astrocytoma because these two histotypes are biologically closer to each other than to oligodendroglioma (40). Furthermore, two cases with tumor percentage less than 50%, determined by DNA (based on ploidy and mutation fraction) and RNA (using the ESTIMATE algorithm) methods, were excluded. Therefore, 23 cases in total were subjected to the ‘response versus transcriptional subtypes’ analysis. The five most effective monotherapies in our drug panel (Figure 3D) were included in this analysis.

A statistically significant positive association was observed between MES gliomas and μCancer response to JQ1 and CC-115, and a similar trend was observed for vorinostat (Figures 5C and 5D). Conversely, a negative correlation was observed between PN gliomas and μCancer response to vorinostat and CC-115, with a similar trend observed for JQ1. These opposing response profiles between PN and MES tumors are consistent with the well-established ‘PN-to-MES switching’ concept that highlights the differences in biological characteristics and therapy responses between these two subtypes (41,42). Similar to MES cases, GPM gliomas exhibited a significant positive correlation with μCancer response to JQ1, vorinostat, and CC-115 (Figure 5D). A trend towards negative correlation was observed between these three agents and NEU gliomas with only vorinostat attaining statistical significance.

A negative correlation was observed for capivasertib μCancer response and CL gliomas (Figure 5C). Intriguingly, navitoclax, which exhibited effective responses across most μCancers, did not show any response preference to the transcription subtypes tested. This suggests that Bcl-2 family proteins represent an important node of vulnerability in the majority of gliomas.

Consistent with the preferential efficacy of tested compounds towards specific transcriptional subtypes, SPL scores alone did not correlate with µCancer response to treatment because exclusively MES, CL, or PN tumors would all be assigned an SPL score of 1 (Figure S5E). However, MES tumors with the highest purity (i.e., SPL close to 1) were most responsive to JQ1, vorinostat, and to a smaller degree CC-115. This supports the positive correlation between MES and μCancer response to these drugs (Figure 5C), although validation in a larger cohort is needed.

Next, we interrogated genes that are differentially expressed between responders and non-responders for drug-subtype pairs that showed significant correlation. By removing cases with responses between responders and non-responders (Figures 4B and S5F), this analysis reduced sample size and statistical power, but allowed identification of genes that differentiate the two groups (Figure 5E; Table S10). For JQ1, the most significantly differentially expressed genes in the MES signature were matrix metalloproteinase MMP7 and glioma pathogenesis-related protein 1 (GLIPR1) (43). For vorinostat, a set of PN-associated genes were downregulated, whereas GPM genes were upregulated in the responder group (Figures 5E and S5G). Of note, μCancers derived from PN gliomas with high expression of ERBB3 appeared to be less responsive to vorinostat. Response to CC-115 correlated with higher expression of MES-associated genes, with extracellular matrix (ECM) remodelers (COL15A1, LUM, COL1A1) topping the list (Figures 5E). Response to the pan-AKT inhibitor capivasertib was negatively correlated to the expression of CL-associated genes (Figure 5E). The data suggest that CL-enriched tumors, which encompass most of the EGFR altered gliomas (38), are less likely to respond to capivasertib.

### Temporal RNA profiling reveals mechanisms of resistance to BET inhibition

To interrogate molecular mechanisms underlying drug response and resistance, we focused on JQ1 for its significant single-agent efficacy. µCancers from a JQ1 responder (PT440) and non-responder (PT431) were selected for RNA-seq analysis following 2 day and 4 day of JQ1 treatment (Figure 6A). We first examined the effect of JQ1 on the temporal expression of key GSC markers (44,45) and transcriptional subtypes, both of which are critical contributors to therapeutic resistance. JQ1 treatment resulted in downregulation of the prominent markers *CD133* and *CD15,* as well as *CD109* in both PT431 and PT440 µCancers, while changes in *ITGA6* and *L1CAM* expression were variable between the two cases (Figure 6A). These data suggest JQ1 can target at least a subset of GSC populations. Notably, JQ1 induced a temporal reduction in MES and its related GPM subtypes, accompanied by a gradual increase in PN enrichment in both µCancers (Figures 6B and S5H). CL subtype enrichment increased during JQ1 treatment only in the responding PT440 µCancers, though levels remained low compared to the non-responding PT431 (Figure 6B). The data suggest that the increased response of PT440 µCancers to JQ1 may relate to their high level of MES enrichment and also points to high initial or drug-induced enrichment in PN and CL cell states as potential mechanisms of JQ1 resistance. These findings further support the opposing roles of MES and PN subtypes in glioma behavior and therapeutic response (41,42). The CL subtype is often associated with activated EGFR signaling (46), and crosstalk between BET and EGFR signaling has been implicated in reciprocal resistance to either monotherapy (47–49). Thus, a durable response to BET inhibition may require an upfront combination or sequential administration of JQ1 followed by an EGFR or other RTK pathway inhibitor, particularly in HGGs with emerging CL enrichment.

**Figure 6.**
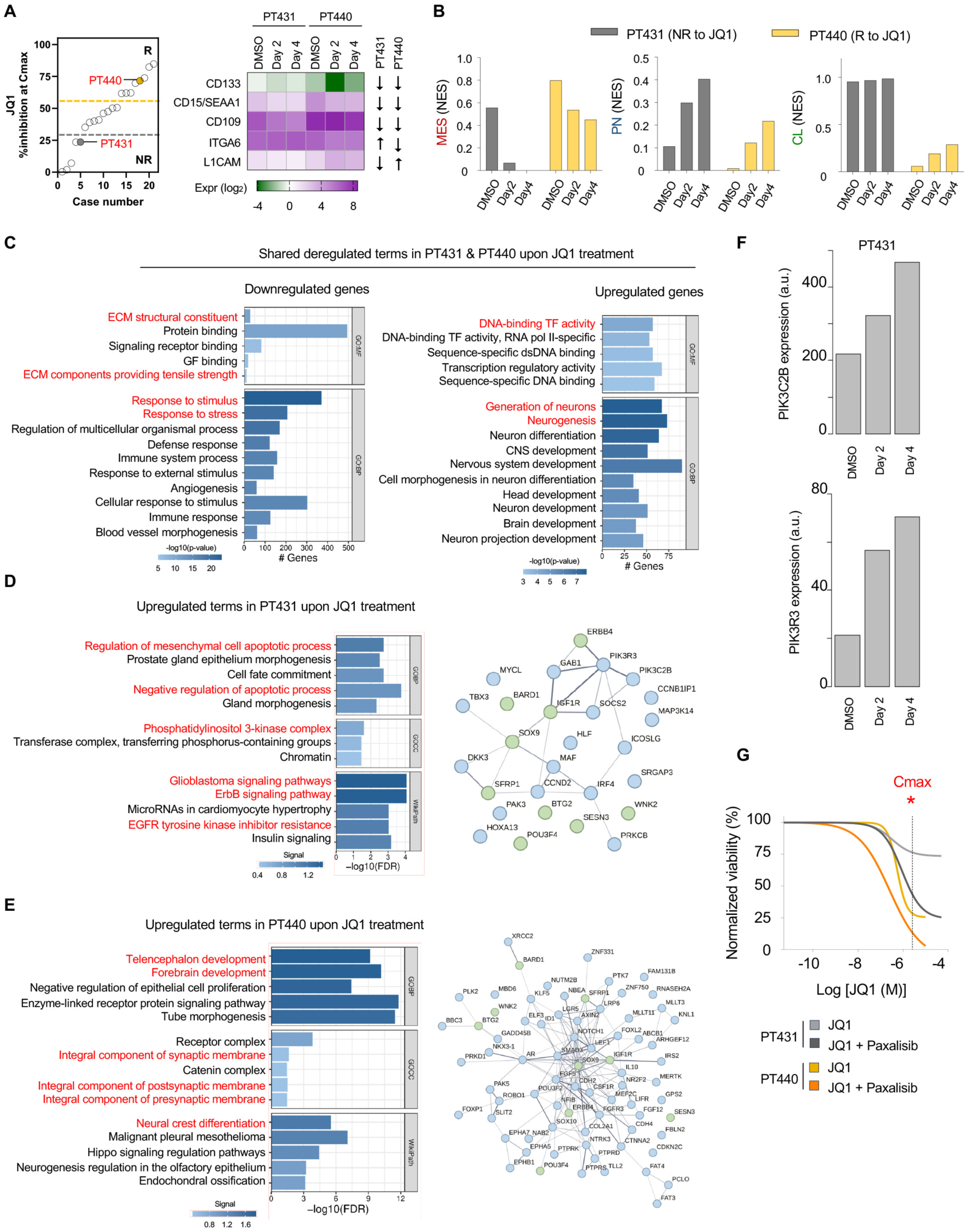
**Molecular mechanisms of μCancer response and resistance to JQ1.** (A) Distribution plots show % inhibition at Cmax (y-axis) from cases with low-to-high response from left to right (x-axis). Responders (R) and non-responders (NR) are cases above the yellow dashed lines and below the grey dashed lines, respectively. Non-responding PT431 and responding PT440 to JQ1 indicated. Heatmap showing expression (Expr; on a natural log (ln) scale) of selected GSC marker upon JQ1 treatment (left), with upregulation (upward arrow) and downregulation (downward) compared to DMSO indicated. (B) Bar graphs showing JQ1-induced temporal changes in the enrichment of TCGA transcriptional subtypes in μCancers. (C) Bar graphs showing JQ1-induced, top-ranked downregulated (left) and upregulated (right) GO terms, shared by PT431 and PT440 µCancers. Pathway analysis based on genes with ≥ 0.1 (log_2_ scale) progressive increase at both day 2 and day 4. (D-E) Bar graphs showing top-ranked upregulated GO terms and WikiPathways (left). Cancer genes with ≥ 0.2 (log_2_ scale) and ≥2-fold increase at day 4 (visualized in protein network, right) selected for pathway analysis for PT431 (D) and PT440 (E). Differentially upregulated (blue) and shared (green) genes between two cases indicated. (F) Bar graphs showing JQ1-induced temporal expression changes of PIK3C2B (top) and PIK3R3 (bottom) in PT431 μCancers. (G) Dose-response curves showing PT431 and PT440 μCancers response to paxalisib alone and in combination with JQ1. Cmax indicated (dashed vertical line topped with asterisk). See also Table S10. Terms in red mentioned in text. Abbreviations: ECM, extracellular matrix; GF, growth factor; TF, transcriptional factor.

Next, we explored shared gene expression changes and pathways in both non-responding PT431 and responding PT440 µCancers, aiming to broadly examine the transcriptional impact of JQ1. ECM remodeling, a process related to mesenchymal behavior (50,51), and response to stress were terms downregulated in both cases (Figure 6C and Table S10). In contrast, shared upregulated gene ontology (GO) terms included transcription regulatory activity, neurogenesis, and CNS development.

To investigate adaptive bypass mechanisms that limit PT431 response to JQ1, we focused on temporally upregulated OncoKB/COSMIC-defined cancer genes for pathway analysis (Figures 6D and 6E). Using the STRING webtool (52) to identify enriched pathways and visualize protein interaction networks, these analyses revealed that upregulation of anti-apoptosis processes and activation of PI3K and RTK pathways may contribute to the limited response of PT431 to JQ1. Indeed, *PIK3C2B* and *PIK3R3,* two positive regulators of PI3K signaling in GBM (53–55) were differentially upregulated by JQ1 in PT431 (Figure 6F). In agreement with this, inhibition of the PI3K pathway with paxalisib markedly increased PT431 µCancer response to JQ1, while having little impact on the already responding PT440 (Figure 6G).

Neurogenesis and synapse-related terms were prominently upregulated in PT440 following JQ1 treatment (Figure 6E) and to a lesser extent in PT431 (Figure 6B). These findings suggest that, similar to radiotherapy which can promote neuron-glioma interactions (56), targeted therapies such as JQ1 may upregulate the neuron-tumor network. Therefore, future strategies combining BET inhibitors with agents that silence neuronal activity or disrupt neuron-tumor connectivity may enhance therapeutic efficacy.

### Combination treatment strategies exhibit strong efficacy across glioma subtypes

Our data argue that combination treatments are more likely to overcome resistance and increase efficacy (57,58). Therefore, we sought to identify drug combination strategies that are more effective than single-agent treatment. Due to the non-proliferative nature of the μcancer assay, which exhausts tumor tissue, we used a rational approach to select agents for combination treatments.

Given the common activation of the RTK pathway in gliomas, we targeted downstream signaling by combining the PI3K/mTOR inhibitor paxalisib with the MEK inhibitor pimasertib. Moreover, by targeting transcriptomic-epigenomic landscapes, epi-drugs can improve sensitivity to other targeted agents (59). Therefore, we included combinations of paxalisib with either vorinostat or JQ1, and pimasertib with JQ1. Given the known functional interactions between BET and HDAC proteins (19), we also included a combination of JQ1 with vorinostat. Further, “apoptosis-inducing agents” (i.e., navitoclax and idasanutlin) can provide therapeutic benefit by triggering cytotoxicity in cancers (60). We thus combined navitoclax with idasanutlin to promote apoptosis by distinct mechanisms, and vorinostat with navitoclax to simultaneously target gene expression and eliminate senescent cells. Overall, seven drug combination treatments were included in our pre-selected drug panel and run in parallel with single agents.

Drug combinations can significantly increase therapeutic benefit without necessarily exhibiting pharmacologically additive or synergistic effects (57,58). Here, we used % inhibition at each drug’s Cmax to compare the effect of single agent treatment versus that of drug combination. In the example provided (Figure 7A, left and middle-left plots), single agent A exhibited 20% inhibition at its Cmax value, while agent B exhibited 62% inhibition at its Cmax. Combination of the two resulted in 77% and 83% inhibition at Cmax for drug A and drug B, respectively. For data visualization, we plotted % inhibition at Cmax for drug A (x-axis) versus that of drug B (y-axis), assigning (20, 62) to single agent effects and (77, 83) to combination treatment. Arrows were then added from single to combination treatment points, with diagonal arrows from left bottom to upper right indicating drug combinations with stronger efficacy than both single agents (Figure 7A, middle-right and right plots). Among seven combinations tested, 6 combination strategies exhibited stronger effects than both of their corresponding single agents, while the combination of navitoclax with idasanutlin was slightly more effective than navitoclax alone (Figure 7B).

**Figure 7.**
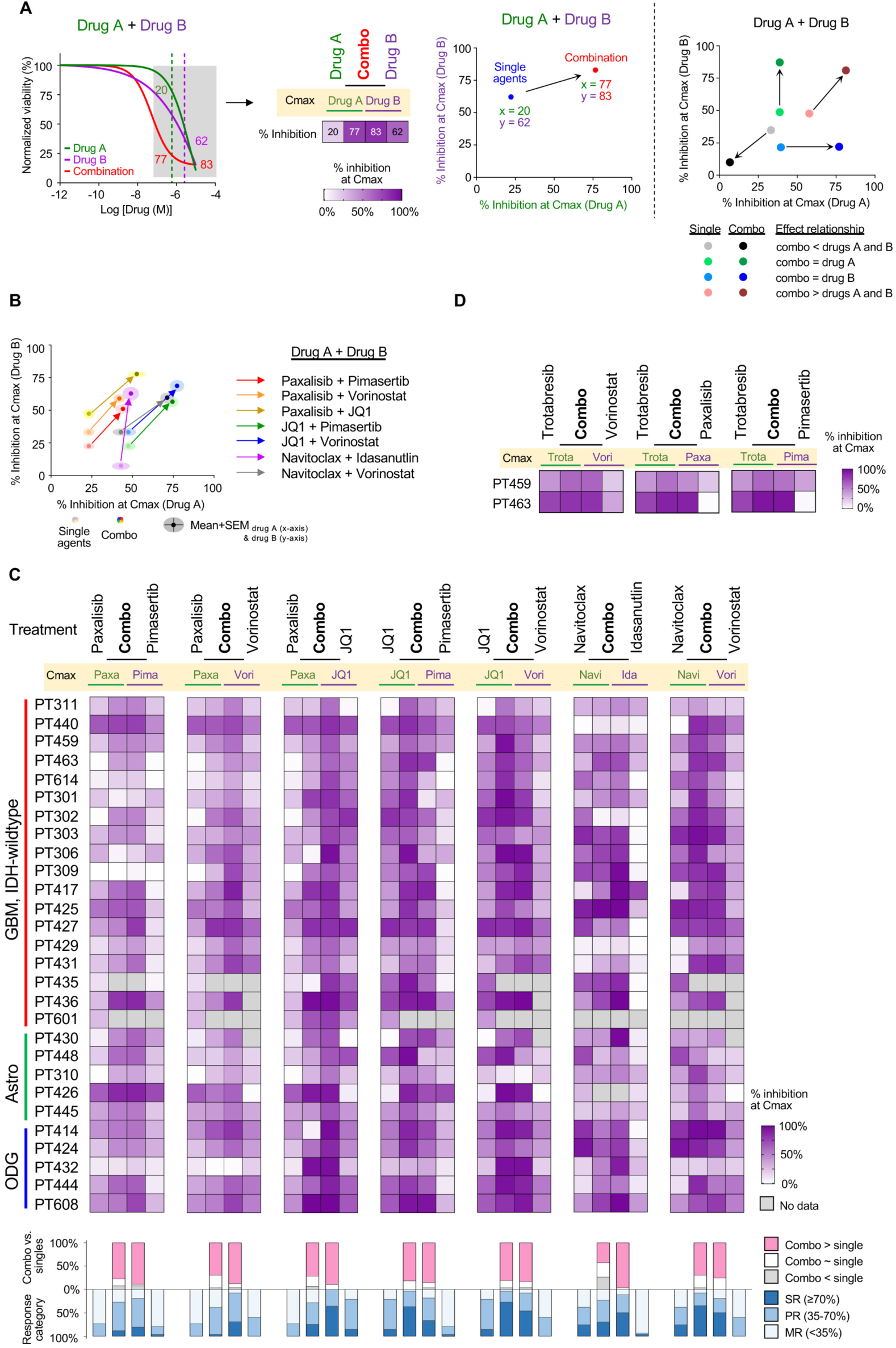
**Effect of combination strategies on μCancer drug responses.** (A) Example representing a dose-response curve graph (left) with indicated % inhibition (grey box) of single agents (drug A, green; drug B, purple) and combination treatment (red) at each drug’s Cmax (vertical dashed lines), conversion to a heatmap (middle-left) or to an X-Y scatter plot (middle-right) showing combination (red) being more effective than corresponding single agents (blue). Potential scenarios (right) of drug combination effects (dark colors) relative to corresponding single agents (light colors) indicated by arrows. (B) Graph depicting the effect of 7 drug combinations (dark colors) compared to their corresponding single agents (light colors). Data presented as mean (solid dots) ±SEM (clouds). (C-D) Heatmaps depicting differential responses of combination treatments compared to single agents at each Cmax. Differential drug response from 0% (white) to 100% (dark purple) indicated. Companion stacked column graph (C) summarizing the relationship between drug combinations and single-agent treatments (top, combinations vs. singles) and response category (bottom) across cases. All drug combinations achieved statistically significant stronger efficacy than single agents as assessed by a multivariable GLMM (*p*<0.0001, see Table S8). See also Tables S7 and S8.

The combination of paxalisib with pimasertib was more effective than single agent treatments in 77% of cases (Figure 7C; Table S7). Combining a molecularly targeted therapy and an epi-drug (i.e., paxalisib plus vorinostat, paxalisib plus JQ1, and pimasertib plus JQ1) similarly resulted in overall stronger effects than monotherapy in 63%, 61%, and 67% of cases, respectively. These results are consistent with the observed upregulation of PI3K and RTK signaling in JQ1 non-responding PT431 μCancers (Figures 6D and 6F) and their role as potential BET inhibitor bypass mechanisms. Combining the two epi-drugs JQ1 and vorinostat was more effective than single agent treatments in 75% of cases. Importantly, we also observed similarly increased efficacy when the clinically relevant BET inhibitor trotabresib was combined with other agents (Figure 7D). Increased efficacy of navitoclax plus idasanutlin was observed in 38% of cases, while combining navitoclax and vorinostat was more effective than single agents in 58% of cases. Importantly, strong responses (≥70% inhibition at Cmax) to combination treatment were observed in 19 to 81% of cases tested (Table S7), with vorinostat plus JQ1 being the most effective strategy. Altogether, the 7 drug combinations produced at least one strong response in 25 out of 28 cases (89%) and were statistically more effective than corresponding single agents (Table S8).

## Discussion

HGGs are aggressive, molecularly complex, and heterogeneous tumors. To explore targeted therapeutic options and identify predictive biomarkers, we employed a functional genomics approach combining genomic-transcriptomic analyses and patient-derived μCancer models. We utilized a model that minimizes selective pressure by not relying on tumor cell expansion and maintains the cellular and molecular diversity of the parental tumor. μCancer growth rates were comparable to that of human tumors, and significantly slower than reported doubling times of conventional models (61). Importantly, μCancers faithfully preserved the genomic and transcriptomic profiles of the original tumors, and their responses to treatments largely recapitulated reported findings in human HGG clinical trials.

Utilizing pre-existing knowledge of oncogenic pathways that drive the growth of HGGs, we developed a glioma-relevant drug panel to interrogate therapeutic vulnerabilities across cases. Agents targeting the PI3K pathway (paxalisib, capivasertib, CC-115), the epigenome (vorinostat, JQ1), and survival/senescence (navitoclax) were more effective than other single agent treatments. RNA expression profiles rather than DNA aberrations were more informative as predictive biomarkers. Finally, highlighting the molecular complexity and heterogeneity of HGGs, combination treatments outperformed single agents and were effective across glioma subtypes.

Despite maintaining focal *EGFR* amplifications, μCancers responded poorly to EGFR-targeted treatments and the broader multi-RTK inhibitors. The data suggest this is a tumor-intrinsic property that also is unrelated to CNS penetration, which is not a barrier for drug efficacy in the μCancer setting. One potential explanation is that RTK amplifications are late events contributing to HGG aggressiveness but are not essential for sustained tumor growth, as also suggested by others (62–64). This hypothesis is further supported by the reported poor efficacies of EGFR/RTK inhibitors osimertinib, neratinib, and sunitinib in glioma clinical studies (Table S5).

Unlike RTK inhibitors, targeting downstream RTK effectors was more effective *ex vivo*, with inhibitors of PI3K signaling exhibiting stronger overall effects than inhibitors of MAPK signaling. Capivasertib induced at least a partial response in 72% of tested cases, CC-115 in 50%, and paxalisib in 27%. Strong responses to treatment were rare. Notably, 26% of patients with progressive or recurrent HGG exhibited metabolic partial responses to paxalisib in a dose-escalation phase I study, while patients with high drug exposure showed significantly longer PFS than the rest of the patient population (Table S5). CC-115 was tested in newly diagnosed *MGMT*-unmethylated GBM patients but was associated with dose-limiting toxicities and was not superior to standard of care in PFS or overall survival (OS) (Table S5). Targeting the DDR pathway with the PARP inhibitor niraparib was ineffective *ex vivo*, which is similar to olaparib in human glioma (65). Finally, drugs targeting epigenetic events, or inducing apoptosis and eliminating senescent cells, were relatively effective in the μCancer setting, with vorinostat inducing at least partial response in 40% of cases, JQ1 in 78% with 4 strong responses, and navitoclax in 62% of cases with 7 strong responses (Table S5). The data argue that strong responses to monotherapy are rare in HGGs but could benefit subgroups of HGG patients.

Notably, we found that RNA expression profiles rather than DNA aberrations were more informative in predicting drug efficacy (Figures 3-6). This is perhaps unsurprising as HGGs are characterized by high molecular complexity and cellular plasticity (36,37). Indeed, transcriptional signatures and pathway enrichments provide more dynamic and integrative readouts of tumor cell behavior, as exemplified by their profound involvement in glioma plasticity (36,37). Accordingly, JQ1 and CC-115 were particularly effective in μCancers enriched for YAP/TAZ activation or derived from MES and GPM gliomas (Figures 4-5). These effects are likely attributed to the role of BET proteins in promoting gene expression and YAP/TAZ activation, as well as DNA-PK in phosphorylating and activating YAP/TAZ (66), collectively inducing YAP/TAZ-mediated transcriptional addiction (31) and an aggressive mesenchymal state (67,68). Finally, μCancers derived from CL gliomas responded poorly to capivasertib, while no correlation was observed between transcriptional subtypes and response to navitoclax.

Using simplicity scoring, we observed that most gliomas in our cohort exhibit multiple transcriptional states. This is in agreement with the known heterogeneity of HGGs (69,70) and the high degree of subclonal events in our tumors. Importantly, our study also highlights the distinct and often opposing therapeutic vulnerabilities between MES and PN or CL subtypes. Specifically, MES-enriched tumors exhibited strong sensitivity to JQ1, vorinostat, and CC-115, whereas PN tumors were poorly responsive (Figure 5C). Providing insight into the mechanism of its action, JQ1 treatment downregulated the MES cell state in both responding and non-responding μCancers while increasing PN and to some extent CL enrichment (Figure 6B). These findings confirm that phenotypic plasticity is a major impediment to effective single-agent treatment and suggest that a combination strategy targeting different subtypes may offer a more effective therapeutic approach for this aggressive disease.

As single agent treatments are unlikely to effectively eliminate such complex tumors, we explored rational combination therapies. These combination strategies exhibited significantly increased efficacy over single-agent treatments, with six strategies being particularly effective in the majority of tested cases (Figure 7 and Tables S7 and S8). Importantly, agents like pimasertib or paxalisib, which showed limited or modest single-agent activity, produced meaningful impact in the combination setting. Combination treatments are also important for epi-drugs, such as BET inhibitors, which are known to succumb to drug resistance (71,72). We observed selective upregulation of anti-apoptosis, EGFR/RTK, and PI3K signaling in JQ1 non-responding PT431 μCancers (Figure 6), suggesting they represent potential mechanisms of drug resistance. This could explain the increased efficacy of JQ1 in combination with the PI3K/mTOR inhibitor paxalisib and the MEK inhibitor pimasertib in our study (Figures 7B and 7C). The increased efficacy of JQ1 in the MES and GPM glioma subtypes also argues that rational combination treatments including clinical BET inhibitors should be explored in the recurrent disease setting, where MES/GPM subtypes are more common and treatment options are ineffective.

Machine learning and artificial intelligence are increasingly been employed to develop models of drug response, primarily trained on large-scale cell line drug-response data (i.e. Cancer Cell Line Encyclopedia pan-cancer data). While the utility of cell line data for HGG drug response has been debated (3), a baseline transformer-based deep learning model that integrates gene expression profiles with drug molecular structure to learn fundamental drug-genome interaction principles could be further fine-tuned by transfer learning using HGG µCancer drug response data. Such a model could potentially predict response to existing therapeutics as well as emerging therapeutics not yet represented in public databases. Alternatively, µCancer pharmacogenomic data could be used to generate multimodal drug prediction models. In both cases, models could be further fine-tuned by actual patient drug response data, as it becomes available from clinical studies, potentially individualizing patient treatment.

In conclusion, this study utilizes a functional genomics approach to identify targeted treatments and predictive biomarkers for HGGs. The data point to select patient populations that could derive clinical benefit from specific drug treatments. They also identify several rational drug combination strategies with strong anti-tumor activity that warrant further pre-clinical and clinical validation. Moreover, further validation and implementation of this approach to the clinic could provide rapid identification of optimal individualized strategies for the treatment of this aggressive disease.

## Methods

### Ethics statement

Patients with HGG who sought consultation from 2019 to 2021 with clinical investigators participating in this study across Mayo Clinic sites were offered participation in the ExVivo study. Patients were enrolled after informed consent under Mayo Clinic Institutional Review Board (IRB)–approved protocol number 13-000942. The study protocol was approved by the research ethics committee IRB of Mayo Clinic. All portions of the study were performed according to the Declaration of Helsinki. All patients provided written informed consent for the use of their tissue, blood, and electronic medical record data.

### Study design

A portion of resected tumor measuring 1-2 cubic cm from each patient was selected by the pathologist and equally split into two pieces. One was immediately flash frozen for nucleotide sequencing and one cryopreserved for μCancer generation and drug screening. Pathological characterization was performed to ensure disease histology. A pre-defined single (*n*=16) and combination (*n*=7) drug screen panel was tested in μCancers of all gliomas with sufficient tissue, except for vorasidenib, which was only tested in IDH-mut cases. The top five most effective agents were subjected to further statistical and bioinformatical analyses for discovery of predictive biomarkers. The increased efficacy of drug combinations was quantified by calculating the differential drug efficacy of combination treatment over that of corresponding single agents at their Cmax. The number of replicates and associated statistical analyses are indicated where appropriate in the figures or figure legends.

### μCancer generation, drug testing, data analysis and quality control

Apportioned fresh tissue specimen was cryopreserved in CryoStor® CS10 cell freezing medium (C2874, Sigma). Thawed tissues were subjected to mechanical and enzymatic dissociation as described previously (11), using Human Tumor Dissociation Kit (130-095-929, Miltenyi Biotec) following the manufacturer’s instructions (Figure 2A). Dissociated cells were resuspended either in: 1) glioma μCancer medium, which contains DMEM/F12 media (SH30023.01, HyClone) supplemented with 10% heat-inactivated horse serum (16050-122, Gibco), 1% penicillin–streptomycin (MT30002CI, Corning), Y-27632 (10 μM; ALX-270-333, Enzo Life Sciences or S1049, Selleckchem), human EGF (10 ng/ml; AF-100-15, PeproTech) and insulin (5 μg/ml; I1882, Sigma); 2) glioblastoma organoid (GBO) medium(73) containing 50% advanced-DMEM:F12 (12634-010, ThermoFisher), 50% Neurobasal (10888022, ThermoFisher), 1X N-2 supplement (17502048, ThermoFisher), 1X B-27 supplement minus vitamin A (12587010, ThermoFisher), 1X GlutaMax (35050-06, ThermoFisher), 1% penicillin–streptomycin (MT30002CI, Corning), 50

μM 2-mercaptoethanol (31350010, ThermoFisher), human insulin (2.5 μg/ml; I1882, Sigma), and Y-27632 (10 μM; ALX-270-333, Enzo Life Sciences); or 3) in neurosphere medium(5) containing Neurobasal (10888022, ThermoFisher) supplemented with 1X N-2 supplement (17502048, ThermoFisher), 1X B-27 supplement minus vitamin A (12587010, ThermoFisher), 1X GlutaMax (35050-06, ThermoFisher), 1% penicillin–streptomycin (MT30002CI, Corning), Y-27632 (10 μM; ALX-270-333, Enzo Life Sciences), EGF (25 ng/ml; AF-100-15, PeproTech), and bFGF (25 ng/ml; AF-100-18B, Peprotech).

An equal number of dissociated cells (5×10^3^) was seeded in 40 μl per well of 96-well Akura^TM^ PLUS hanging drop plates (CS-03-001-00, InSphero) and cultured for 6 days allowing for μCancer generation. Established μCancers were then transferred to the corresponding well of 96-well clear round bottom ultra-low attachment microplates (7007, Corning) and subjected to microscopic evaluation to ensure size similarity across wells, before drug testing at eight serial dilutions with corresponding drug solvents (DMSO or media) as vehicle controls. After 6-days of drug treatment, cell viability was quantified using the CellTiter-Glo luminescent assay (G7570, Promega). The luminescence output was interpolated and transformed into nM ATP by comparison to an ATP standard curve. Data were then transformed into Log10, and the dose-response curves were fitted on the basis of four parameter logistic regression with occasional use of three-parameter logistic regression for cases with insufficient points to establish a top and bottom plateau. Upward curves were deemed 0% growth inhibition. Cases with inconsistent μCancer sizes and low ATP values (<10 nM) and poor curve fit were deemed quality control failures and excluded from subsequent analysis.

Drug efficacy was represented as percentage of reduced viability at Cmax compared to the baseline vehicle control. For the pink stacked bar graph listed below the heatmap in Figure 7C, drug combinations were considered more effective when yielding ≥9.5% increased efficacy than both corresponding single agents.

### Sequencing pipeline and bioinformatic processing

MPseq, WES, and RNAseq were performed after isolation of DNA and RNA from flash-frozen tumor tissue and analyzed as before (11). Specifically, tumor was identified by gross and frozen section microscopic examination in the Mayo Clinic Frozen Section Laboratory. Fresh tumor in excess of diagnostic clinical material was placed in a plastic cassette and snap frozen in isopentane.

A secondary review by an experienced surgical pathologist was performed using a toluidine blue-stained frozen section. A tumor cellularity of >40% in the sample was obtained that included the use of macrodissection or laser capture microdissection (74) as needed. DNA and RNA were extracted with the Qiagen AllPrep DNA/RNA mini kit (80204, Qiagen) or Qiagen DNeasy Blood and Tissue Kit (69504, Qiagen) following the manufacturer’s protocol. In cases where laser capture microdissection was done or initial DNA yield was too low for sequencing (PT427), whole genome amplification was performed as previously described (74) using the Repli-G mini kit (150023, Qiagen).

Whole genome sequencing was performed on all tumor tissue samples using Nextera Mate-Pair Kit (FC-132-1001, Illumina) and MPseq, a low-pass whole-genome DNA sequencing method. All libraries were sequenced at an average depth of 71.5 bridged-coverage, sufficient for DNA junction detection. The sequencing fragment data were mapped to reference genome GRCh38 by BIMA (75) with junctions and CNAs called by SVAtools (76). Junctions were detected from clusters of discordant (based on the mapped location and/or orientation of the reads in each fragment) fragments spanning two breakpoints (76). CNAs were called based on read-depth of concordant fragments (77).

CNAs were determined by the comparison with the genome ploidy for each case (Table S3). Amplification was called when a significant copy number gain was observed (e.g., ≥7n for a diploid genome). Molecular alterations were deemed pathogenic when TSGs exhibited LoF mutation(s) or complete inactivation, or when oncogenes exhibited GoF mutation(s), copy number amplification, and/or oncogenic fusion transcripts, as reported before (2).

µCancers from select cases (PT301, PT427, PT431, PT440, PT459) were also collected for RNAseq. Cells were grown in hanging drop plates (5×10^3^ cells per well) for 6 days for µCancer establishment, µCancers collected at indicated times, and used for RNA purification (QIAGEN RNeasy Micro Kit, Cl) and human mRNA sequencing using the NovaSeq X Plus Series (PE150) platform. For all 5 cases, established non-treated µCancers from one to two 96-well hanging drop plates were collected. For PT431 and PT440, µCancers were also transferred to ultra-low attachment plates and treated with JQ1 or DMSO for 2 days or 4 days. One to three 96-well plates for each condition were collected for RNAseq.

### RNA landscape comparison between μCancers and corresponding patient tumors

To determine the consistency in gene expressions between each µCancer and its corresponding patient tumor, RNA-seq expression summaries (“count” files) from both types of samples (tumor tissue, n=30; µCancer treatments, n=17) were processed together in R. A group testing analysis using lmFit and eBayes functions in limma package was employed to eliminate batch effects that might obscure true biological differences, between the two groups. Following, a dendrogram including patient tumors and μCancers was generated by hclust function with “maximum” distance method and “complete” clustering method. Additionally, to determine gene expression similarities, log normalized gene expression data was generated separately for patient tumors and μCancers and used to plot and estimated their correlations between the two types of samples.

### Transcription signature subtyping

RNA-seq data from patient tumors and μCancers were processed through our MAP-Rseq (78) and Novagene pipelines, respectively to generate gene expression “.count” files. These files were then input in the edgeR package in R to generate log normalized gene expression values. Normalized enrichment score (NES) for each sample was obtained by the “gsva” and “ssgseaParam” functions in GSVA R package. To determine the subtype classification of a tumor, the NES for gene sets based on either TCGA (38) or biological pathways (39) were calculated and scaled to a range between zero and one and then the subtype with the highest score was assigned to that tumor. The “simplicity score” was computed as the difference between the highest NES and the average of the NES in other subtypes (38). The function ‘geneData’ in the ‘gage’ R package was used to generate heatmaps of differentially expressed genes (*p*<0.05). Finally, Sankey diagrams were generated by the “sankey” package in R. To determine the effects of drugs on the subtype character of μCancers, enrichment scores on day 2 and day 4 after therapy were compared against DMSO treatments.

### Temporal changes of gene expression and pathway enrichment upon drug treatment

To determine the dynamics of drug response and resistance, we utilized two related gene set analysis tools. First, genes were selected for downstream g:Profiler pathway analysis (79) if they had a minimum of 0.1 (log_2_ scale) progressive change at both day 2 (vs. DMSO) and day 4 (vs. day 2) of JQ1 treatment in both non-responding PT431 and responding PT440 µCancers. In the second analysis, we identified genes with a minimum of 0.2 (log_2_ scale) increase at day 2 vs. DMSO and at day 4 vs. day 2 in PT431 and PT440 µCancers. Also, we required a minimum of 2-fold increase on day 4 compared with DMSO. Selected genes were then additionally filtered for cancer genes in the OncoKB and COSMIC databases. Finally, filtered genes were input into the STRING database to identify enriched gene sets in each of the two JQ1-treated µCancers. Bar graphs were generated using the SRplot webtool (80).

**Differential ploidy analysis.** By using a recently developed algorithm (81), we examined correlations between drug responses and an integrated copy number score termed here as differential ploidy which was calculated as a continuous variable based on copy number changes of genes in the targeted signaling pathways. Our algorithm corrected for the confounding effects due to high ploidy including whole genome doubling (WGD) which is commonly seen in advanced tumors, including HGGs (82–84). Without this correction, CNA estimations are often erroneously biased towards copy-number gains in high-ploidy samples. After correcting for high ploidy, a score for a given drug-targeted signaling pathway from KEGG database was generated for each tumor using cumulative aberration of CNAs for each gene relative to the corrected ploidy value. Additionally, our algorithm accounted for oncogenes in the gained areas and TSGs in the deleted regions by incremental increases in differential ploidy scores. On the other hand, differentially ploidy scores were incrementally decreased for any loss of known oncogenes or gain of known tumor suppressor genes.

### *pTERT* genotyping by PCR

The *TERT* promoter sequence at the two known pathogenic sites C228 and C250 was amplified with M13-TERT genotype primers (forward: 5’-GTAAAACGACGGCCAGACGTGGCGGAGGGACTG-3’ and reverse: 5’-CAGGAAACAGCTATGACAGGGCTTCCCACGTGCG-3’) using Q5® high-fidelity DNA polymerase (M0492S, NEB) (85). Amplified PCR segments were then gel purified for Sanger sequencing using standard M13 sequencing forward and reverse primers (Genewiz).

### Ki-67 staining

Paraffin-embedded GBM8 xenograft tissue expressing NT-sh virus (86) and GBM8 μCancers along with PT425 tumor tissue and resulting PT425 μCancers in 5 μm sections were stained for human Ki-67 (M7240, Dako) following standard autostainer (Link 48, Dako) protocol. Images were acquired using the Aperio AT2 slide scanner (Leica Biosystems).

### Cell lysate preparation, antibodies, and Western blotting

Briefly, µCancer spheres from two 96-well plates were collected and lysed in 100 µl of RIPA buffer (50 mM Tris-HCL, pH 7.4, 150 mM NaCl, 1% NP-40, 0.5% NaDC, 0.1% SDS) containing a proteinase inhibitor (P50700, RPI) and a phosphatase inhibitor (78427, ThermoFisher). The spheres were then subjected to needle shearing (29G for 7 times) with vortexing to maximize cell lysis. 5-10 µg of protein lysates were resolved by SDS-PAGE, and Western blotting was performed using ECL (45-00-875, ThermoFisher) and PICO (34580, ThermoFisher) detection reagents. Antibodies used in this study included p70 S6 kinase (9202, CST), p-Thr389 p70 S6 kinase (9205, CST), GAPDH (2118, CST), and HRP-conjugated anti-rabbit secondary antibody (711-035-152, Jackson ImmunoResearch).

### Statistical analysis

Drug data processing and statistical analysis were performed using the GraphPad Prism software. Unpaired t-test was performed when assessing the impact of *p*TERT status on μCancer response to JQ1. Correlation analysis was performed using the Spearman and/or Pearson methods as indicated in figure legends. Astrocytomas and GBMs with ≥50% of tumor purity (exclusion of PT417 and PT429) were included for statistical correlation analyses. Tumor purity was inferred based on DNA content (mutation allele fraction and ploidy) and/or RNA expression data (ESTIMATE algorithm (87)). Correlation analysis was performed using the “cor.test” function in R software package with Pearson’s correlation coefficient outputs. Statistical significance was assessed at *p*-value <0.05.

A multivariable, generalized linear mixed model (GLMM) was constructed to evaluate the response (i.e., percent inhibition at Cmax) of *ex vivo* µCancers to treatment with drug given alone or in combination. The statistical model included fixed effects for the drug, combination versus monotherapy administration, and the interaction of the drug and whether it was given in combination. Random (indirect) effects were modeled to account for 1) within-patient variability (i.e., patient sample), and 2) residuals for the random effect of repeated drug regimens, with a compound symmetry covariance structure using SAS v9.4 (Cary, NC). Finally, a supplemental model was constructed to assess the direct effect of distinct regimens while accounting for within-patient variability as above.

## Data availability

All data associated with this study are included in this published article or its in supplementary information. Additional information is available from the corresponding authors upon request, following the institution’s policy and regulation.

## Acknowledgments

The authors thank the patients and their families for donating their specimens to make this study possible and providing hope for future patients with this devastating disease. We also greatly appreciate the support from the clinical staff and translational study team. Regretfully, we lost one of our team members, Giannoula Karagouga, who has significantly contributed to this work, to a car accident during manuscript preparation. We appreciate Dr. Ken Buetow (Arizona State Univeresity) and Dr. Jann Sarkaria and Will Herbert (Mayo Clinic Rochester) for helpful discussions as well as reagents from Dr. Sarkaria. We also thank Dr. Alan R. Penheiter (Mayo Clinic Rochester) and Dr. Mark A. Edgar (Mayo Clinic Florida) for their assistance in validating somatic variants and performing Ki-67 analysis on patient tissue, respectively. This work was supported by the Mayo Clinic Center for Individualized Medicine and its Biomarker Discovery and Precision Cancer Therapeutics Programs. Additionally, PZA was supported by National Institutes of Health grant P30 CA15083 and National Institute of Neurological Disorders and Stroke grant R01 NS101721, and SHK was supported by the U.S. Food and Drug Administration R01 FD-R-07288.

## Author contributions

Conceptualization: ASM, MJB, JCC, GV, PZA

Methodology: WHL, FK, JBS, RWF, SHJ, SJM, JCC, GV, PZA

Investigation: WHL, FK, JBS, MTB, RWF, JH, DS, SS, SHJ, FRH, TB, AFM, SJM, LK, LEH, DMG, SFE, LY, CMI, MAS, WJS, ABP, SSR, SHK, KAJ, MMM, TCB, JCC, GV, PZA

Visualization: WHL, FK, JBS, SHJ, SJM Funding acquisition: ASM, MJB, JCC, GV, PZA Project administration: ARE, LJ, JSK

Supervision: SHK, KAJ, MMM, ASM, MJB, BRB, TCB, AQH, JCC, GV, PZA

Writing – original draft: WHL, PZA Writing – review & editing: All authors

## Supplementary information

Document S1. Figures S1-S5, Tables S1-S10, and supplementary references S1 – S40.

