## Supplementary data for "Functional genomics identifies therapeutic options, biomarkers, and resistance mechanisms for high-grade gliomas"

### Supplementary information

Figures S1-S5, Tables S1-S10, and supplementary references S1 – S40.

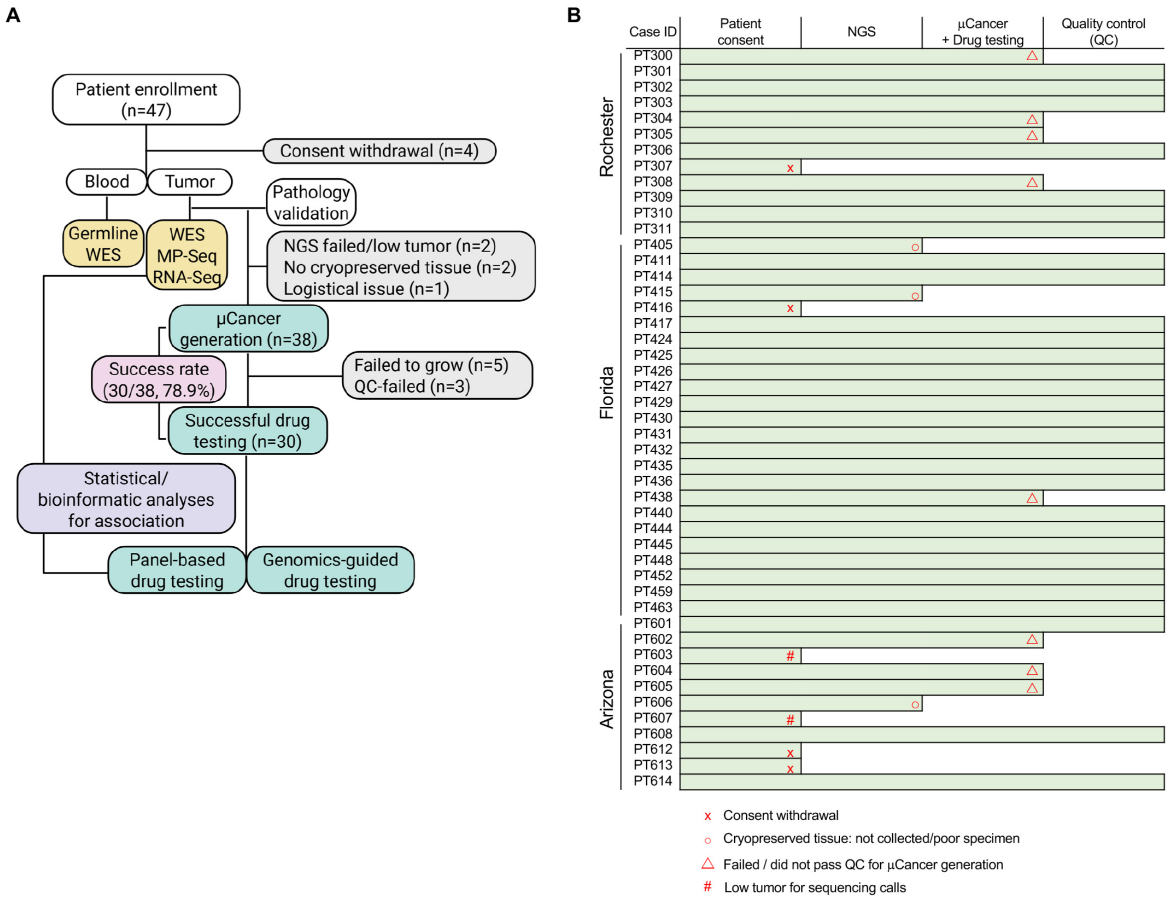


**Figure S1. Overall study and individual case workflow.**

(A) Flow chart indicating the number of patients enrolled and proceeded with following applications for NGS (yellow) and µCancer generation and drug testing (green). Success rate of µCancer establishment (pink) and reasons for non-evaluability (gray) indicated.

(B) Trajectory of each case from patient enrollment, next-generation sequencing (NGS), µCancer generation and drug testing to quality control.

Related to Figure 1A.

**Figure S2**

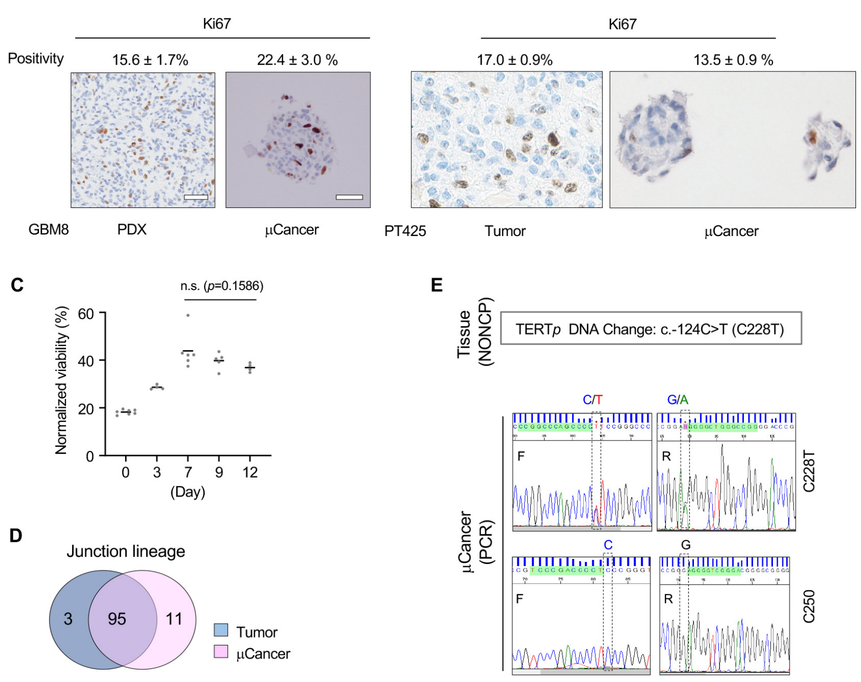


**Figure S2. μCancer characterization and comparison with parental GBM.**

(A-B) Representative images of PDX GBM8 (A) and PT425 (B) tumor tissue (left) and corresponding μCancers (right) at 7 and 6 days *ex vivo,* respectively, stained for Ki-67 proliferation marker. Combined percentage (mean±SE) of Ki-67 positive cells from multiple images indicated (n=3-6). Scale, 50 mm.

(C) μCancer (PT425) cell viability measured by nM ATP over 12-day of culture in at least quadruplicate with mean value indicated (line).

(D) Venn diagram illustrating the similarity and difference of DNA junction numbers captured by MPseq in μCancer (pink) and corresponding parental tumor PT311 (blue).

(E) *pTERT* mutation status in the parental tumor PT311 (top) and μCancers (bottom) identified by clinical testing (Neuro-Oncology expanded gene panel (NONCP), top) and PCR (bottom), respectively. Sanger sequencing output from forward (bottom, F) and reverse (bottom, R) primers at pathogenic *pTERT* mutation regions (C228 and C250) are shown. Heterozygous C228T and unaltered C250 indicated.

### Figure S3

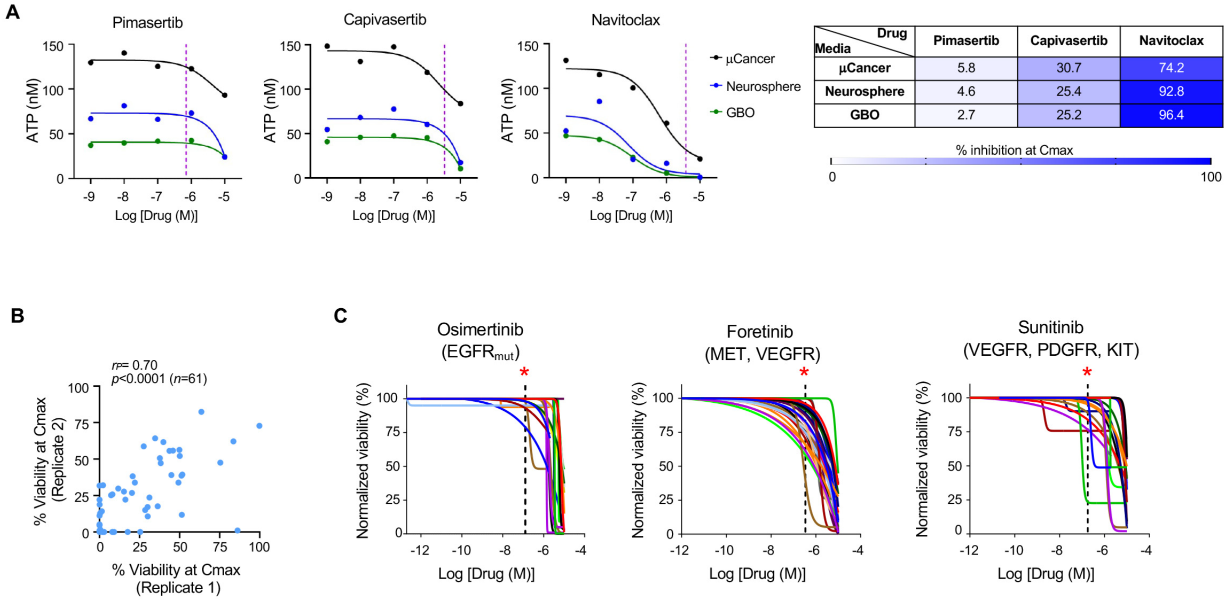


**Figure S3. μCancer culture conditions and drug response comparison.**
(A) Dose-response curves of μCancers (PT303) treated with indicated agents when grown in indicated media (μCancer: black; neurosphere, blue; glioblastoma organoid (GBO), green). Cmax values are indicated (pink vertical dashed lines). Heatmap shows % inhibition at Cmax for each condition (far right).
(B) Scatter plot depicts % viability of all replicates performed at different times of drug testing (x-axis, 1st experiment; y-axis, 2nd experiment), showing correlation between experiments. Each data point represents drug responses measured at the 1st and 2nd experiments. Pearson's ($r_p$) correlation analysis performed.
(C) Normalized dose-response curves for indicated agents targeting indicated RTKs. Cmax indicated (vertical dashed lines with asterisks above).
Related to Figure 3.

**Figure S4**

A

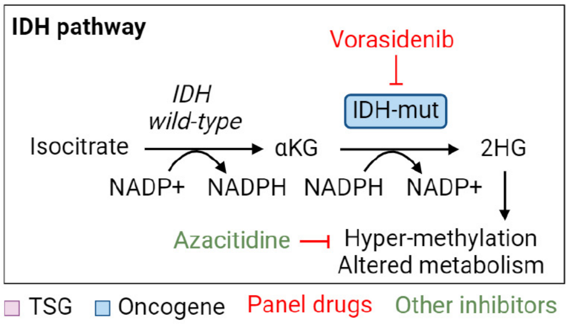


B

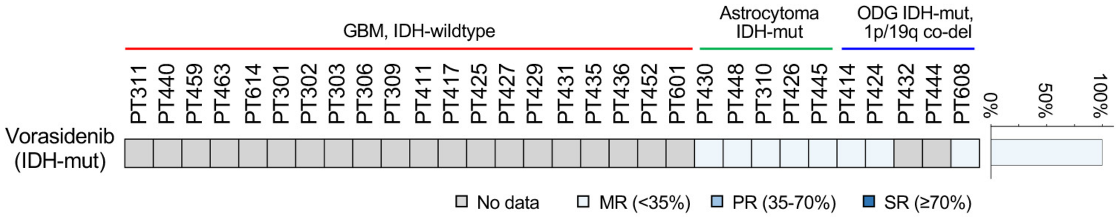


**Figure S4. Ex vivo response of IDH-mut µCancers to vorasidenib monotherapy.**

(A) Schematic illustrating the IDH signaling pathway and potential targeted strategies.

(B) Three-tier drug response heatmap showing <35% (minimal response, MR; light blue), 35-70% (partial response, PR; blue), and ≥70% (strong response, SR; dark blue) of growth inhibition at Cmax for vorasidenib. Stacked bar graph (right) showing the percentage of differential responses.

See also Table S6.

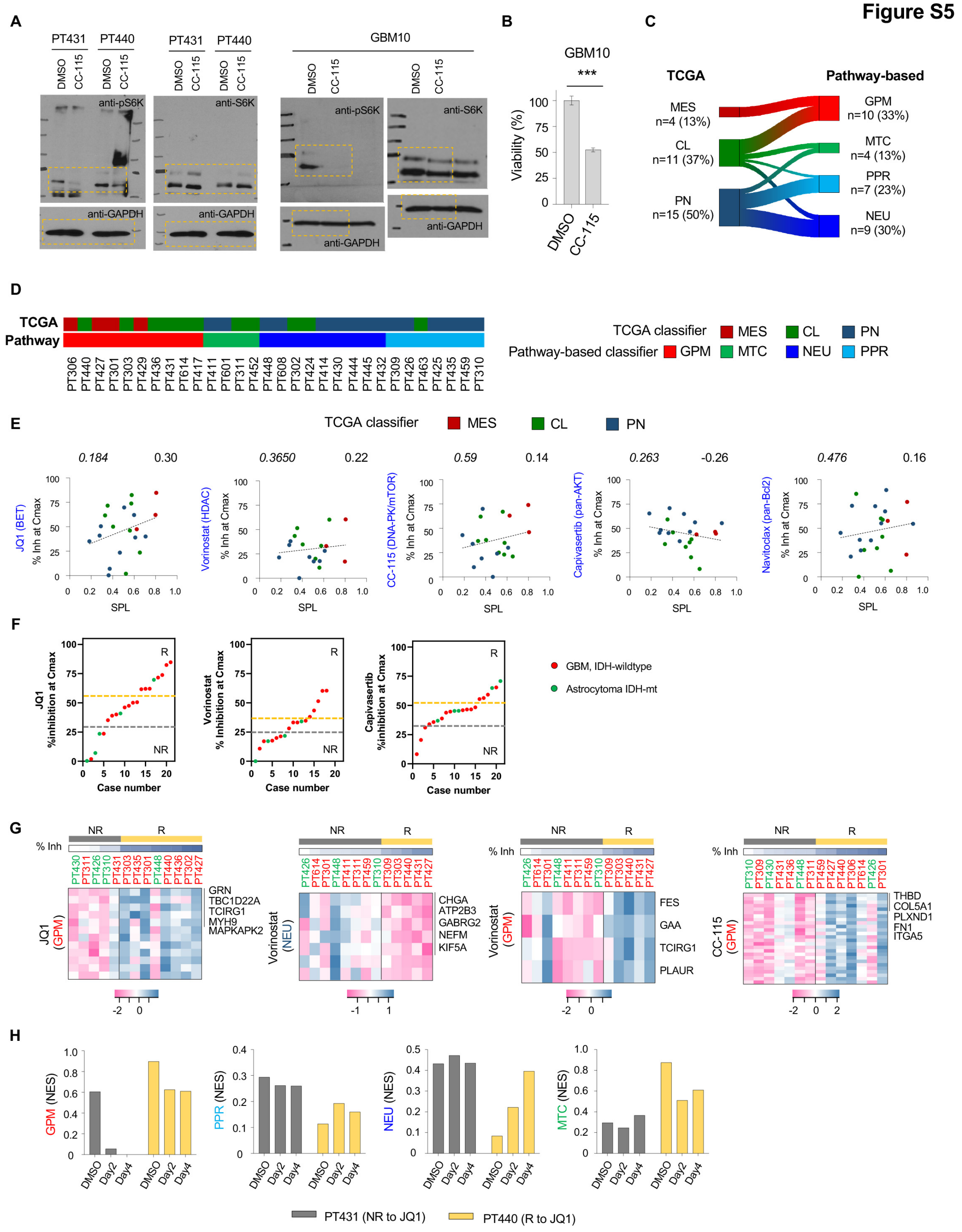


**Figure S5. Transcription classification underlying µCancer therapeutic vulnerability.**
(A) Uncropped gel images for boxed regions shown in Figure 4B.

(B) Bar graph showing relative viability (mean±SEM) of GBM10 μCancers treated with CC-115 at Cmax vs. DMSO control (***$p$<0.0005, unpaired t-test).
(C) Sankey diagram depicting the association of assigned subtypes between the TCGA (left) and pathway-based (right) transcription classifiers in glioma. Number and percent of cases in the cohort indicated.
(D) Subtype assignment using the TCGA (top) and pathway-based (bottom) transcription classifications for individual cases.
(E) Correlation plots showing the relationship between the SPL scores (x-axis) and μCancers response to indicated agents (y-axis). Pearson's correlation coefficient (upper right) with p-values (upper left) indicated. Histotypes for each case color-coded.
(F) Distribution plots show % inhibition at Cmax (y-axis) from cases (red dots, GBM; green dots, astrocytoma) with low-to-high response from left to right (x-axis). Responders (R) and non-responders (NR) are cases above the yellow dashed lines and below the grey dashed lines, respectively.
(G) Heatmaps showing differentially expressed genes for indicated gene sets between treatment responder (R, yellow) and non-responder (NR, gray) μCancer groups. Z-scores from low (dark magenta) to high (dark cerulean) and % inhibition at Cmax (% Inh) indicated. Top 5 differentially expressed genes listed.
(H) Bar graphs showing temporal changes of pathway-based transcriptional subtypes in μCancers upon JQ1 treatment.

**Table S1. Baseline demographic and clinical data of patients included in the drug screen.**

**Table S1**

| | Glioblastoma, IDH-wildtype (n=20) | Astrocytoma, IDH-mutant (n=5) | Oligodendroglioma, IDH-mutant and 1p/19q co-deleted (n=5) |
|---|---|---|---|
| Disease status, *n* (%) | | | |
| Newly diagnosis | 5 (25) | 2 (40) | 1 (20) |
| Recurrent / progressing | 15 (75) | 3 (60) | 4 (80) |
| Age at enrollment (years) | | | |
| Median (range) | 58 (28-85) | 33 (29-46) | 51 (31-63) |
| Sex, *n* (%) | | | |
| Male | 6 (30) | 5 (100) | 5 (100) |
| Female | 14 (70) | 0 (0) | 0 (0) |
| Previous treatment (n, %) | | | |
| No treatment | 5 (25) | 2 (40) | 1 (20) |
| Radiation | 14 (70) | 3 (60) | 4 (80) |
| Temozolomide | 14 (70) | 3 (60) | 3 (60) |
| Nitrourea-based therapy | 4 (20) | 0 | 1 (20) |
| Bevacizumab | 5 (25) | 0 | 1 (20) |
| Tumor treating fields | 2 (10) | 0 | 0 |
| Others | 3 (15) | 0 | 0 |
| Previous treatment lines (n, %) | | | |
| 0 | 5 (25) | 2 (40) | 1 (20) |
| 1 | 10 (50) | 3 (60) | 1 (20) |
| 2 | 3 (15) | 0 | 2 (40) |
| 3+ | 2 (10) | 0 | 1 (20) |
| MGMT promoter status, n (%) | | | |
| Methylated | 9 (45) | 2 (40) | 2 (40) |
| Unmethylated | 9 (45) | 1 (20) | 0 |
| Indeterminant | 1 (5) | 1 (20) | 0 |
| Unknown | 1 (5) | 1 (20) | 3 (60) |
| pTERT status, n (%) | | | |
| C228T | 11 (55) | 0 | 4 (80) |
| C250T | 5 (25) | 0 | 1 (20) |
| Non-C228T/C250T (WT or VUS) | 2 (10) | 4 (80) | 0 |
| Indeterminant | 1 (5) | 0 | 0 |
| Unknown | 1 (5) | 1 (20) | 0 |

**Table S2.** (See Excel spreadsheet)

**Table S2. Demographic and clinical characteristics of the ExVivo-glioma cohort.** Notes: Tempus, Tempus xT gene panel; pMGMT clinical tests performed before (* ) or after ( *) the ExVivo-glioma study indicated. Disease diagnosis and corresponding 2016 and/or 2021 WHO classification indicated whenever necessary. For newly diagnosed cases, see Diagnosis at the ExVivo-glioma period for initial diagnosis. Related to Figure 1 and Table 1.

Abbreviations: n/a, not applicable; NONCP, neuro-oncology expanded gene panel; PCR, polymerase chain reaction; pMGMT, MGMT promoter methylation status; RT, radiation therapy; TMZ, temozolomide; TTF, tumor treating fields; VUS, variant of unknown significance.

**Table S3.** (See Excel spreadsheet)

**Table S3. Genomic alteration information to complement Figure 1B.** Note: Accrual of pathogenic alterations for the histogram in Figure 1B highlighted.

Abbreviations: amp, amplification; cnLOH, copy-neutral loss of heterozygosity; ESS, essential splice site; FS, frameshift; GoF, gain of function; LB, likely benign; LoF, loss of function; SG, stop-gained; VUS, variant of unknown significance.

**Table S4. Full gene list identified by WES for case PT311.** Related to Figure 2.

| | | Allele fraction | |
|---|---|---|---|
| **Gene** | **Alteration** | **Tissue** | **μCancer** |
| PTEN | c.395G>T_p.Gly132Val | 0.70 | 0.52 |
| OR7D2 | c.608C>T_p.Thr203Met | 0.56 | 0.62 |
| NPPA | c.427G>A_p.Gly143Arg | 0.48 | 0.41 |
| VPS13D | c.9769G>A_p.Val3257Ile | 0.45 | 0.27 |
| OLIG2 | c.339C>G_p.Ile113Met | 0.45 | 0.39 |
| RPAP1 | c.1039C>T_p.Arg347Trp | 0.43 | 0.39 |
| EYS | c.8654G>C_p.Gly2885Ala | 0.41 | 0.38 |
| SHISA2 | c.25G>A_p.Val9Ile | 0.40 | 0.46 |
| COL24A1 | c.274A>G_p.Thr92Ala | 0.40 | 0.26 |
| ZADH2 | c.781G>T_p.Ala261Ser | 0.40 | 0.26 |
| FAM83H | c.1216G>A_p.Ala406Thr | 0.39 | 0.37 |
| IL31RA | c.1150G>C_p.Val384Leu | 0.39 | 0.41 |
| MYO5B | c.2486G>A_p.Arg829His | 0.38 | 0.39 |
| WDR49 | c.829C>T_p.Arg277Trp | 0.37 | 0.37 |
| MIA3 | c.4408G>T_p.Asp1470Tyr | 0.37 | 0.36 |
| CAPN9 | c.205C>T_p.Arg69X | 0.37 | 0.38 |
| DNAH17 | c.12299G>A_p.Arg4100His | 0.36 | 0.44 |
| CEP170B | c.3785G>A_p.Arg1262Gln | 0.36 | 0.43 |
| TBC1D14 | c.137A>T_p.Tyr46Phe | 0.35 | 0.013 |
| SLC25A48 | c.623C>G_p.Ser208Cys | 0.35 | 0.37 |
| DSG3 | c.2117C>T_p.Thr706Met | 0.34 | 0.32 |
| NR5A2 | c.485G>A_p.Arg162His | 0.33 | 0.24 |
| ELFN1 | c.1951G>A_p.Ala651Thr | 0.32 | 0.30 |
| MYH1 | c.5257C>T_p.Arg1753Cys | 0.32 | 0.43 |
| SIGLEC1 | c.2819C>A_p.Ser940X | 0.31 | 0.20 |
| PALM | c.575C>T_p.Thr192Met | 0.28 | 0.27 |
| RASAL3 | c.1907G>A_p.Arg636His | 0.25 | 0.22 |
| ANO2 | c.2909G>A_p.Arg970Gln | 0.24 | 0.34 |
| SLC1A6 | c.169G>A_p.Ala57Thr | absent | 0.21 |

**Table S5** (also provided in Excel spreadsheet)

| Drug | Target | CNS penetration | FDA approved indications | IC50 (nM) | Cmax µM (Log M) | Latest development phase for glioma | Key clinical trial/ case report findings |
|---|---|---|---|---|---|---|---|
| **Abemaciclib** | CDK4/6 | Yes<br>Ms[S1]<br>Hs[S2] | Breast Cancer (early stage and metastatic/advanced disease) | 2 (CDK4)*#<br>10 (CDK6)*# | 0.312<br>(-6.506)[S2] | Phase 2 | NCT02981940[S3]<br>- Patient population: rGBM exhibiting loss of CDKN2A/B with intact RB<br>- Treatment: abmeaciclib<br>- Result: 1) From 31 evaluable patients, best response include PR (n=1, 3.2%), SD (n=11, 35.5%) and PD (n=19, 61.3%) with the PFS6 rate being 9.37% (95% CI, 2.4% to 22.27%).<br><br>NCT02977780 (INSIGhT)[S4]<br>- Patient population: ndGBM with unmethylated MGMT<br>- Treatment: Adjuvant abemaciclib following concurrent TMZ/RT<br>- Results: 1) Patients from the abemaciclib arm (6.3 months) showed improved mPFS (HR, 0.72; 95% CI, 0.49 to 1.06; one-sided *P*=0.046, long-rank test) compared with the control arm (4.6 months); 2) No significant improvement in OS was found; 3) No improved PFS in the abemaciclib arm for CDK-positive patients. |
| **Vorinostat** | HDAC | Yes<br>Ms[S5]<br>Hs[S6] | Cutaneous T-cell lymphoma | 10 (HDAC1)*#<br>30 (HDAC3)* | 1.2<br>(-5.921)†§ | Phase 2 | NCT00238303[S6]<br>- Patient population: rGBM received one or fewer chemotherapy for progressive disease<br>- Treatment: Vorinostat monotherapy<br>- Results: A slightly progression-free benefit was found in patients treated with vorinostat dose escalated to 300 mg for two or more cycles, warrnting future exploration for vorinostat in combination strategies.<br><br>NCT00731731 (Alliance N0874/ABTC 02)[S7]<br>- Patient population: ndGBM<br>- Treatment: Vorinostat in combination with SoC<br>- Result: There was no improvement in OS.<br><br>NCT00939991[S8]<br>- Patient population: rGBM<br>- Treatment: Vorinostat in combination with bevacizumab and daily TMZ<br>- Results: PFS6 was not statistically improved compared to historical controls. |
| **Neratinib** | pan-ERBB (EGFR ERBB2 ERBB4) | Yes<br>Ms[S9]<br>Hs[S4] | Breast Cancer (early stage and metastatic disease) | 92 (EGFR)*#v<br>59 (ERBB2)*#v<br>19 (ERBB4)v | 0.146<br>(-6.836)[S10] | Phase 2 | NCT02977780 (INSIGhT)[S4]<br>- Patient population: ndGBM with unmethylated MGMT<br>- Treatment: adjuvant neratinib following concurrent TMZ/RT<br>- Result: 1) statistically significantly improved mPFS in the neratinib arm (6.1 months) compared to the control arm (4.7 months) (HR, 0.72; 95% CI, 0.50 to 1.02; one-sided *P*=0.033, long-rank test); 2) No significant improvement was found in OS; 3) EGFR-enriched subpopulation showed significantly longer PFS but not OS compared with the control group (HR, 0.42; 95% CI, 0.24 to 0.72; one-sided *P*< 0.001, long-rank test). |
| **Osimertinib** | EGFR-mut | Yes<br>Ms[S11]<br>Hs[S12] | EGFR-mut NSCLC (early-stage, CNS metastasis) | 1<br>(L858R/T790M)*<br>12 (L858R)* | 0.126<br>(-6.900)[S13] | Phase 2 | Case series[S14]<br>- Patient population: rGBM with simultaneous EGFR ampliciation and EGFRvIII (n=15)<br>- Treatment: bevacizumab with off-label osimertinib<br>- Results from retrospective analyses: Osimertinib plus bevacizumab combination was marginal effective (mPFS, 5.1 months (95% CI 2.8.-7.3); mOS, 9.0 months (95% CI 3.9-14.0)) in this patient population with some experiencing a long-lasting clinical benefit. |
| **Sunitinib** | RTK (VEGFRs, PDGFRs, KIT) | Unclear**<br>Ms[S15]<br>Hs: case report[S16] | Gastrointestinal stromal tumor<br><br>Pancreatic neuroendocrine tumor<br><br>Renal cell carcinoma (advanced/high risk of recurrent) | 80 (VEGFR2)<br>2 (PDGFRβ) | 0.181<br>(-6.7423)[S13] | Phase 2 | NCT00535379 (SURGE 01-07)[S17]<br>- Patient population: rGBM (first recurrence)<br>- Treatment: Sunitinib<br>- Results: 1) Minimal anti-tumor activity was found (PFS6, 12.5%; mPFS, 2.2 months; mOS, 9.2 months); 2) Patients with high expression of c-KIT in vascular endothelial cells have better PFS.<br><br>[S18]<br>- Patient population: rHGG (WHO grade III/IV supratentorial glioma)<br>- Treatment: Sunitinib<br>- Results: 1) None of the patients achieved an objective respose. 2) There was no correlation between treatment reponse and copy numbers/protein expression of sunitinib targets (KDR, PDGFRA, KIT). |
| **Pimasertib** | MEK | Yes<br>Ms[S19] | no | 5 (MEK1/2)# | 0.7<br>(6.155)[S19] | Preclinical | n/a |

| | | | | | | | |
|---|---|---|---|---|---|---|---|
| **Paxalisib** | PI3K<br>mTOR | Yes<br>Ms[S20]<br>Hs[S21] | no | 2 (PI3Kα)[#]<br>46 (PI3Kβ)[#]<br>3 (PI3Kδ)[#]<br>10 (PI3Kδ)[#]<br>70 (mTOR)[#] | 0.58<br>(-6.237)[§] | Phase 2 | NCT01547546[S22]<br>- Patient population: r-HGG (WHO grade III/IV)<br>- Treatment: Paxalisib<br>- Results: Patients with higher exposure to paxalisib (Cmax and AUC estimated by MR-PET imaging) have significantly improved PFS compared with patients with lower drug exposure (HR=0.4176; 95% CI, 0.1821-0.9576; *P*=0.0039 and HR = 0.4679; 95% CI, 0.1801-1.216; *P*=0.0296, respectively).<br><br>NCT01547546[S21]<br>- Patient population: progression or recurrent HGG<br>- Paxalisib<br>- Results: The MTD for paxalisib was 45 mg/day; at this dose, metabolic partial response determined by FDG-PET was observed in 26% (7 of 27) patients.<br><br>NCT03522298[S23]<br>- Patient population: ndGBM with unmethylated MGMT<br>- Treatment: Paxalisib<br>- Results: Paxalisib at MTD showed prolonged PFS and improved OS. |
| **Capivasertib** | AKT | Unknown | Breast Cancer (HR+/HER2- locally advanced/ metastatic with PIK3CA/AKT1/ PTEN alterations; with fulvestrant) | 3 (Akt1)*[#]<br>7-8 (Akt2)*[#]<br>7-8 (Akt3)*[#] | 3.32<br>(-5.479)[S24] | Preclinical | n/a |
| **Idasanutlin** | MDM2 | Yes<br>Ms[S25] | no | 6*[#] | 6.08<br>(-5.216)[S26] | Phase 1/2<br>(Recruiting) | n/a |
| **Ulixertinib** | ERK | Yes<br>Ms[S27]<br>Hs: case report[S28] | no | <3 (ERK1/2)*[#] | 6.34<br>(-5.198)[S28] | Phase 1<br>(Recruiting) | n/a |
| **Foretinib** | RTK<br>(MET, VEGFR2) | Yes<br>Ms[S29] | no | 0.4 (Met)*[#]<br>0.9 (VEGFR2)*[#] | 0.34<br>(-6.469)[S30] | Preclinical | n/a |
| **Navitoclax** | pan-Bcl2<br>(Bcl-2, Bcl-xL, Bcl-w) | Yes<br>Ms[S31] | no | ≤0.5 (Bcl-xL)[#]<br>≤1 (Bcl-2)[#]<br>≤1 (Bcl-w)[#] | 3.8<br>(-5.420)[S32] | Preclinical | n/a |
| **Niraparib** | PARP | Yes<br>Ms[S33] | Ovarian cancer (recurrent/advanced) | 3.8 (PARP1)*[#]<br>2.1 (PARP2)*[#] | 2.5<br>(-5.602)[†] | Phase 2 | NCT05076513[S34]<br>- Patient population: ndGBM<br>- Treatment: Niraparib<br>- Results (preliminary): PFS6 was 64% (n=11) at a median follow-up of 8.1 months (6-12.9 months). |
| **JQ1** | BET<br>(BRD1, BRD2, BRD4) | Yes<br>Ms[S35] | no | 77 (first bromodomain, BRD1/2/4)*[#]<br>33 (second BD, BRD1/2/4)*[#] | 2.58<br>(-5.588)[S36] | Preclinical | n/a |
| **CC-115** | DNA-PK<br>mTOR | Yes<br>Hs[S37] | no | 13 (DNA-PK)*[#]<br>21 (mTOR)*[#] | 0.223<br>(-6.652)[S37] | Phase 2 | NCT02977780 (INSIGhT)[S4]<br>- Patient population: ndGBM with unmethylated MGMT<br>- Treatment: CC-115 concurrent with RT and adjuvant monotherapy<br>- Results: CC-115 showed was associated with high toxicity and no PFS/OS benefit was found. |
| **Vorasidenib** | IDH-mut | Yes<br>Hs[S38] | no | 0.04-22<br>($IDH1_{mut}$)*<br>7-14 ($IDH2_{mut}$)* | 0.47<br>(-6.328)[S39] | Phase 3 | NCT04164901 (INDIGO)[S40]<br>- Patient population: r/r-LGG (WHO grade 2) with IDH mutations who are treatment naive<br>- Treatment: Vorasidenib<br>- Results: mPFS was significantly improved in the vorasidenib arm (27.7 months; 95% CI, 17.0 to not estimated) compared to the placebo group (11.1 months; 95% CI, 11.0 to 13.7) (HR for progression or death, 0.39; 95% CI, 0.27 - 0.56; *P*<0.001) |

**Table S5. Pharmacokinetic measures and key clinical trial findings of panel drugs used in the study.** Notes: Cmax information was obtained from human clinical trials except JQ1, †DailyMed (https://dailymed.nlm.nih.gov/dailymed/), or §manufacturer. Clinical trial identifiers (NCT#) and trial names indicated where available. CNS penetration property was determined based on CNS response data from mouse (Ms) and human (Hs) studies, and/or pharmacokinetics data where available. IC50 was obtained from drug vendors (MedChemExpress* and Selleckchem#). **, pharmacokinetics data from mouse studies showed limited penetration of the BBB; however, CNS efficacy in a case of human renal cell carcinoma was reported. Supporting evidence included.

Abbreviations: CI, confidence interval; FDG-PET, fluorodeoxyglucose-positron emission tomography; HR, hormone receptor; mPFS, median progression-free survival; MR-PET, magnetic resonance/positron emission tomography; MTD, maximum tolerated dose; ndGBM, newly diagnosed glioblastoma; OS, overall survival; PCV, procarbazine, lomustine and vincristine; PD, progressive disease; PFS6, 6-month progression-free survival; PR, partial response; rGBM, recurrent glioblastoma; rHGG, recurrent high-grade glioma; r/r-LGG, residual or recurrent low-grade glioma; RT, radiation therapy; SD, stable disease; SoC, standard of care; TMZ, temozolomide; TTF, tumor-treating fields; n/a, not applicable.

**Table S6.** (See Excel spreadsheet)

**Table S6. Percent inhibition at Cmax of tested single agents across cases.** Related to Figures 3D and 3E.

**Table S7.** (See Excel spreadsheet)

**Table S7. Percent inhibition at Cmax of tested drug combinations across cases.** Note: data presented in Figure 7 highlighted in yellow.

**Table S8.**

**Table S8. Drug efficacy comparison using multivariable, generalized linear mixed-effects modeling.**

**Table S9.** (provided in Excel spreadsheet)

**Table S9. Gene lists to complement heatmaps in Figure 4A.**

**Table S10.** (provided in Excel spreadsheet)

**Table S10. Gene lists to complement heatmaps in Figures 4C, 5E, S5F, and 6C-6E.**
